# Cardiolipin remodeling maintains the inner mitochondrial membrane in cells with saturated lipidomes

**DOI:** 10.1101/2024.05.29.596368

**Authors:** Kailash Venkatraman, Itay Budin

## Abstract

Cardiolipin (CL) is a unique, four-chain phospholipid synthesized in the inner mitochondrial membrane (IMM). The acyl chain composition of CL is regulated through a remodeling pathway, whose loss causes mitochondrial dysfunction in Barth syndrome. Yeast has been used extensively as a model system to characterize CL metabolism, but mutants lacking its two remodeling enzymes, Cld1p and Taz1p, have not recapitulated the structural and respiratory phenotypes observed in other systems. Here we show the essential role of CL remodeling in the structure and function of the IMM in yeast grown under reduced oxygenation. Microaerobic fermentation, which mimics natural yeast environments, caused the accumulation of saturated fatty acids and, under these conditions, remodeling mutants showed a loss of IMM ultrastructure. We extended this observation to HEK293 cells, where iPLA_2_ inhibition by bromoenol lactone resulted in respiratory dysfunction and cristae loss upon mild treatment with exogenous saturated fatty acids. In microaerobic yeast, remodeling mutants accumulated unremodeled, saturated CL, but also displayed reduced total CL levels, highlighting the interplay between saturation and CL biosynthesis and breakdown. We identified the mitochondrial phospholipase A_1_ Ddl1p as a regulator of CL levels, and those of its precursors phosphatidylglycerol and phosphatidic acid, under these conditions. Loss of *DDL1* partially rescued IMM structure in cells unable to initiate CL remodeling and had differing lipidomic effects depending on oxygenation. These results introduce a revised yeast model for investigating CL remodeling and suggest that its structural functions are dependent on the overall lipid environment in the mitochondrion.

## Introduction

Cardiolipin (CL) is a mitochondrially-synthesized phospholipid (PL) whose structure contains two phosphates and four acyl chains. In eukaryotes, CL is synthesized from the condensation of mitochondrial phosphatidylglycerol (PG) and cytidine diphosphate diacylglycerol (CDP-DAG), both of which are derived from phosphatidic acid (PA) that is imported from the ER (1). CL makes specific interactions with electron transport chain (ETC) proteins and their supercomplexes, which are required for optimal function of the ETC and ATP synthesis (2–5). Depending on its chemical environment, CL can behave as a non-bilayer lipid due to its negative spontaneous curvature (6). Through this property, CL could also function in shaping the morphology of the IMM (7), particularly its highly curved cristae membranes, and in mitochondrial fission-fusion dynamics (8–12).

The acyl chain composition of CL is determined post-synthetically through a remodeling pathway. In humans, heart CL is typically tetra linoleic (18:2) while in the brain and gut there is a wider distribution of CL acyl chain identities, with oleic (18:1) chains as the most common (13–15). Despite this relative heterogeneity, the most abundant form of CL in every organism contains unsaturations on each acyl chain (14, 16). CL acyl chain composition results from the coordinated action of two types of remodeling enzymes, phospholipases and a transacylase, responsible for converting immature CL, which contains saturated chains, to a mature, tetra-unsaturated form of CL (17–20). Phospholipase A_2_ (iPLA_2_) enzymes first initiate CL remodeling by deacylation of nascent CL into monolysocardiolipin (MLCL) (21, 22). The transacylase Tafazzin (TAZ) then catalyzes the reacylation of MLCL into a homogeneously unsaturated CL by transferring unsaturated acyl chains from mitochondrial PLs (17, 23). Abnormal CL compositions have been linked to human pathologies and are thought to drive mitochondrial dysfunction (24, 25).

CL remodeling is lost in patients with Barth syndrome (BTHS), an X-linked disease which arises from a mutation in the *TAZ* gene (26, 27). BTHS patients have altered CL profiles, characterized by a reduction in CL, but an increase in MLCL and more saturated CL (28–31). This altered CL metabolism is thought to cause the mitochondrial defects that result in cardiac and skeletal muscle myopathies as well as neutropenia, muscle weakness and fatigue (32, 33). Numerous studies have specifically implicated the large increase in the MLCL:CL ratio as the major cause of mitochondrial dysfunction observed in BTHS (34–36), and BTHS diagnosis is often performed by assessment of this ratio (37). Identification of the negative impacts of MLCL in mitochondrial membranes has led to therapeutic efforts to prevent its formation. This has been achieved through treatment with bromoenol lactone (BEL), an iPLA_2_ inhibitor (38). BEL mitigates the MLCL:CL ratio in patient lymphoblasts and stabilizes ETC complexes in HEK293 models of BTHS (21, 39). However, despite the reduction in MLCL, BEL-treated cells do not recover respiratory function to control levels (40). Thus, it remains unclear if MLCL is the only molecular driver of mitochondrial dysfunction in BTHS. Notably, BEL-treated cells still accumulate unremodeled CL (22), whose contribution to respiratory and morphological defects have yet to be shown in mitochondria independently of MLCL.

Budding yeast (*Saccharomyces cerevisiae*) has been used extensively as a model system to investigate CL remodeling and cellular mechanisms underlying BTHS (41, 42). While mammals contain multiple CL remodeling processes (24), yeast only perform Tafazzin-mediated remodeling. Unlike mammalian cells, yeast knockouts in the CL synthesis and remodeling pathway are viable, allowing for their phenotypic dissection. In yeast, a single phospholipase A_2_ (Cld1p), conducts the deacylation of nascent CL to MLCL (43). Tafazzin (Taz1p) then transfers an unsaturated acyl chain from mitochondrial PC to CL to make tetra oleoyl (18:1) cardiolipin (TOCL) (14, 44, 45), the maximum level of unsaturation as *S. cerevisiae* are unable to synthesize polyunsaturated fatty acids (PUFAs). Deletion of the *TAZ1* gene results in increased MLCL, reduced CL and more saturated CL species (35, 46, 47)*. CLD1* knockouts also lack remodeled CL but exhibit a milder reduction in its abundance and increased saturation (43, 47). Previous reports have shown that *taz1*Δ, but not *cld1*Δ, cells display temperature-sensitive respiratory growth phenotypes (46, 48, 49), increased oxidative stress (50), and defective bioenergetic coupling (51). These phenotypes are less severe than those occurring from the complete loss of CL in *crd1*Δ (52). Despite these observations, it has been surprising that these mutants do not possess altered mitochondrial morphology or ultrastructure (47). The combined *cld1*Δ*taz1*Δ strain contains an identical PL profile to *cld1*Δ, extinguishes the MLCL:CL ratio, and has been shown to rescue respiratory defects in *taz1*Δ (47, 49, 53).

CL is just one of several bulk PLs in the IMM and its functional roles could be dependent on its local lipid environment. In a previous study, we demonstrated that CL synthesis was essential for the formation of highly-curved cristae in the IMM under conditions of increased lipid saturation induced by genetic or environmental repression of the acyl-CoA desaturase Ole1p (54). Oxygen is a substrate of the Ole1p reaction and is bound to the desaturase through a weak, di-iron binding site (55, 56); microaerobic growth thus acts as a natural modulator of lipid saturation. Saturation modulates the extent of ATP synthase oligomerization at cristae ridges, which CL mechanically buffers against (54).

Here we report phenotypes for loss of CL remodeling in yeast cells with saturated lipidomes. We show that microaerobic growth causes CL remodeling mutants to show a characteristic loss of cristae membranes in yeast mitochondria, which were previously only observed in mammalian models of BTHS. We observe a stronger dependence of IMM ultrastructure on Cld1p activity compared to Taz1p, suggesting that accumulation of unremodeled CL is more deleterious than MLCL for the structure of the yeast IMM in this condition. In mammalian cells, we show that iPLA_2_ inhibition only results in mitochondrial abnormalities when co-treated with the saturated fatty acid, palmitate (Palm). We find that mitochondrial defects in *cld1*Δ yeast cells correspond to a loss of total CL and are partially rescued by deletion of the gene encoding for the mitochondrial phospholipase Ddl1p, which has wide-ranging effects on the composition of CL and its anionic PL precursors.

## Materials and Methods

*Yeast strains and growth conditions*: All *S. cerevisiae* strains used in this study are described in Table S1. For aerobic experiments, cells were grown in complete supplement mixture (CSM) (0.5% Ammonium Sulfate, 0.17% yeast nitrogen base without amino acids and 2% glucose) lacking appropriate amino acids for selection. Yeast mutants were generated by PCR-based homologous recombination, ORFs were replaced by either KanMX (*TAZ1, DDL1)*, *TRP1 (CLD1) or HIS3 (ATG32),* except for *ddl1*Δ*taz1*Δ which was generated by replacing the *DDL1* ORF with KanMX and the *TAZ1* ORF with *HIS3*.

For aerobic growth conditions, cells were grown overnight in a controlled temperature shaker (200 rpm) at 30°C in CSM synthetic media containing 2% glucose using 14 mL culture tubes (18 mm diameter) with a snap/vent cap (Greiner). Cells were then backdiluted into 5 mL fresh CSM and then grown until stationary phase prior to analysis. For microaerobic growth, cells were first pre-incubated in CSM containing 2% glucose in a controlled temperature shaker overnight at 30°C. Aerobic overnight cultures were then backdiluted 1:25 into 24 mL fresh CSM media (25 mL total volume) in 27 mL elongated glass culture tubes (Pyrex) and subsequently fitted with 15 mm diameter rubber stopper caps with holes, which were connected via tubing to a water source to allow gas outflow while limiting inflow. The tubes were placed in a 30°C incubator (without shaking) and grown for 48 hours. After growth, cells reached an OD_600_ of ∼0.8-1.0. Subsequent live cell imaging was conducted on aliquots, and cells were lysed and flash frozen for lipidomic analysis.

*Mammalian cell culture*: HEK293 cells (Sigma-Aldrich) were cultured in DMEM (Gibco) supplemented with 10% FBS (Gibco) in 37°C incubators containing 5% CO_2_. For bromoenol lactone (BEL) treatment, cells were seeded and grown for 48 hours in 2.5 μM BEL (Cayman Chemical), 100 μM palmitate (Palm) or both, prior to analysis. Palm was prepared as previously described (57): Sodium palmitate (Thermo Fisher) was complexed to fatty acid free BSA. Palm was dissolved 1:1 in a mixture of 0.1 mM NaOH in HBSS and ethanol to a concentration of 200 mM, heated to 65°C, and then added to a 4.4% BSA HBSS solution that had been heated to 37°C such that the final Palm concentration was 2 mM in a 3:1 molar ratio with BSA. The mixture was then stirred vigorously and incubated at 37°C for 1 hour. The Palm:BSA mixture or 4.4% BSA vehicular control was diluted into complete DMEM containing 10% FBS to a concentration of 100 μM and filtered through a 2 μm PES filter (Fisher Scientific) before use. Cells were seeded and then grown with complete DMEM containing Palm complexed to BSA and subjected to analysis after 48 hours. The treatment time was selected as there are no observable defects to organelle morphology, but an increase in esterified saturated fatty acids (57). Cells were routinely tested for mycoplasma and not detected.

*Confocal microscopy*: Live cell microscopy was conducted using Plan-Apochromat 63x/1.4 Oil DIC M27 objective on the Zeiss LSM 880 with an Airyscan detector; image acquisition and processing was performed with ZEN software using default processing settings. To assess mitochondrial morphology in yeast grown in either aerobic or microaerobic conditions, strains were transformed with IMM-localized Cox4-GFP and cells were subsequently grown in CSM containing 2% glucose aerobically overnight prior to back dilution into fresh media and incubation in aerobic or microaerobic growth conditions. After 24 hours (aerobic) or 48 hours (microaerobic) of incubation, cells were imaged in 8-well coverglass-bottom chambers (Nunc Lab-Tek) pre-treated with concanavalin A (Sigma Aldrich). Cells were imaged using a 488 nm laser line. N>50 cells in biological triplicates from each condition were then grouped into either ‘normal’, encompassing interconnected morphologies, and ‘abnormal’, consisting of fragmented and completely ablated mitochondrial structures, as has been previously described (54).

For analysis of mitochondrial membrane potential in yeast strains, aliquots of aerobic or microaerobic cells were incubated with 200 nM of membrane potential sensitive dye tetramethylrhodamine ethyl ester (TMRE, Thermo Fisher Scientific T669). Cells were first incubated in the dark with dye for 20 minutes at room temperature followed by three washes in water prior to imaging with a 561 nm laser line. Relative levels of TMRE fluorescence were quantified in the midplane of N=20 cells in each condition using imageJ. Analysis of mitochondrial nucleoids was performed by staining cells with SYBR Green I (SGI, Thermo Fisher Scientific S7563). Cells were first washed with PBS prior to staining with SGI (1:10,000) for 10 minutes at room temperature in the dark. After staining, cells were washed 3 times with PBS prior to imaging using a 488 nm laser line at 0.010% laser power. The number of nucleoids per cell were quantified in N>50 cells. Finally, relative levels of reactive oxygen species (ROS) were determined in each strain in aerobic or microaerobic conditions by staining with 20 μM of ROS-sensitive dye (H_2_DCFDA, Thermo Fisher Scientific) for 30 minutes covered from light. Cells were then washed 3 times with PBS prior to imaging using a 488 nm laser line. DCFDA intensity was quantified from N=20 cells in each condition. For each image, a subsequent DIC image was taken to ensure that dead cells were not included in the quantification. For staining of vacuole membranes, cells were grown in either CSM containing 2% glucose until stationary phase (aerobic), for 48 hours (microaerobic) or for 72 hours in CSM containing 3% glycerol. Cells were then stained with 10 μM FM4-64 (MedChemExpress) for 30 minutes in a 30°C shaker in the dark prior to washing once with media. Cells were then resuspended in fresh media and incubated in a 30°C shaker in the dark for 90 minutes. After incubation, cells were washed twice with media prior to imaging using a 561 nm laser line.

For HEK293 samples, cells were first seeded into individual 35 mm Mattek dishes and subjected to treatment with either 100 μM PA, 2.5 μM BEL or a combination of both, for 48 hours prior to staining with 200 nM Mitotracker Deep Red (Thermo Fisher Scientific) for 30 minutes. Cells were then washed three times with HBSS, resuspended in clear DMEM without phenol red, and imaged using 633 nm excitation at 0.2% laser power.

*Lipidomics analysis*: Mass spectrometry-based lipid analysis was performed by Lipotype GmbH (Dresden, Germany) as previously described (58, 59). Lipids were extracted using a chloroform/methanol procedure (60). Samples were spiked with internal lipid standard mixture containing: cardiolipin 14:0/14:0/14:0/14:0 (CL), ceramide 18:1;2/17:0 (Cer), diacylglycerol 17:0/17:0 (DAG), lyso-phosphatidate 17:0 (LPA), lyso-phosphatidyl-choline 12:0 (LPC), lyso-phosphatidylethanolamine 17:1 (LPE), lyso-phosphatidylinositol 17:1 (LPI), lyso-phosphatidylserine 17:1 (LPS), phosphatidate 17:0/14:1 (PA), phosphatidylcholine 17:0/14:1 (PC), phosphatidylethanolamine 17:0/14:1 (PE), phosphatidylglycerol 17:0/14:1 (PG), phosphatidylinositol 17:0/14:1 (PI), phosphatidylserine 17:0/14:1 (PS), ergosterol ester 13:0 (EE), triacylglycerol 17:0/17:0/17:0 (TAG). After extraction, the organic phase was transferred to an infusion plate and dried in a speed vacuum concentrator. The dry extract was re-suspended in 7.5 mM ammonium formiate in chloroform/methanol/propanol (1:2:4, V:V:V). All liquid handling steps were performed using Hamilton Robotics STARlet robotic platform with the Anti Droplet Control feature for organic solvents pipetting.

Samples were analyzed by direct infusion on a QExactive mass spectrometer (Thermo Scientific) equipped with a TriVersa NanoMate ion source (Advion Biosciences). Samples were analyzed in both positive and negative ion modes with a resolution of R_m/z=200_=280000 for MS and R_m/z=200_=17500 for MSMS experiments, in a single acquisition. MSMS was triggered by an inclusion list encompassing corresponding MS mass ranges scanned in 1 Da increments (61). Both MS and MSMS data were combined to monitor EE, DAG and TAG ions as ammonium adducts; LPC and PC as an formiate adduct; and CL, LPS, PA, PE, PG, PI and PS as deprotonated anions. MS only was used to monitor LPA, LPE and LPI as deprotonated anions; Cer as formiate adduct.

Data were analyzed with in-house developed lipid identification software based on LipidXplorer (62, 63). Data post-processing and normalization were performed using an in-house developed data management system. Only lipid identifications with a signal-to-noise ratio >5, and a signal intensity 5-fold higher than in corresponding blank samples were considered for further data analysis.

*Respirometry*: For analysis of respiration in HEK293 cells, cells were treated for 48 hours prior to analysis on the Agilent Seahorse XF pro (Agilent). 15000 cells were seeded in biological replicates (n=4) into 96 well seahorse cell culture microplates (Agilent) pre-treated with fibronectin (5 μg/mL). Samples were analyzed using the Seahorse Cell Mito Stress Test, with sequential addition of Oligomycin (1.5 μM), FCCP (1 μM) and a Rotenone/Antimycin A mixture (0.5 μM). Analysis was performed on respiration rates after treatments with inhibitors to readout basal, ATP-linked, maximal and proton leak respiration rates. After the assay, cells were trypsinized and mixed with trypan blue and cell-counted to normalize respiration rates.

*Electron microscopy*: After growth in CSM with 2% glucose, aerobic or microaerobic yeast cells were filtered and fixed in a 3% glutaraldehyde solution for 1 hour at room temperature and then at 4°C overnight. Fixed samples were then treated with 0.25 mg/mL zymolyase 20T for 1 hour at room temperature prior to washing twice with 0.1M sodium cacodylate buffer. Samples were then embedded prior to imaging using a JEOL 1400 transmission electron microscope. HEK293 cells were first cultured to confluency in each treatment condition prior to fixation in pre-warmed 4% glutaraldehyde solution for 30 minutes at room temperature. Samples were then re-fixed and embedded prior to imaging using a JEOL 1400 transmission electron microscope.

IMOD and ImageJ software were utilized to measure cristae tubule lengths and OMM areas from thin section images. For morphological quantification, N=40 mitochondria were counted in each condition and assigned either tubular, shortened or flat/empty structural features. Flat/empty structures included mitochondria bereft of cristae as well as those with IMM structures that were adjacent to the OMM. Cristae lengths were determined from N=30 mitochondria in each condition. For analysis of cristae density in HEK293 cells, N=30 mitochondria were analyzed.

## Results

### Microaerobic growth conditions promote IMM biogenesis and saturated lipidomes

Native yeast environments, like rotting fruit or fermentation vessels, feature lower oxygen levels compared to those often utilized in laboratory studies. In mammalian cells, reduction of oxygen results in hypoxic stress that renders significant changes to mitochondrial structure, reducing elongation (64). We were thus surprised to observe that yeast grown in microaerobic conditions featured mitochondria with elongated cristae, in contrast to shorter structures commonly seen in aerobically-grown cells (Figure 1A). These elongated cristae are more likely to be sheet-like, with greater resemblance to mammalian mitochondria (Figure 1B), while shorter cristae in aerobic cells likely correspond to tubular structures as has been previously described (65). Cristae elongation could be ascribed to increased expression of genes involved in mitochondrial biogenesis and ETC complexes, but analysis of RNA-seq data previously acquired (66) showed decreased expression of all major ETC components and ATP synthase subunits under microaerobic conditions (Figure 1C). Instead, increases in microaerobic expression were observed for the CL remodeling genes, *CLD1* and *TAZ1*, suggesting a heightened role for this pathway under low oxygen (Figure 1C). Changes in expression of CL remodeling enzymes was not observed for CL synthase itself (*CRD1*) and were exclusive to cells subjected to extended long-term microaerobic growth, rather than short-term hypoxia (Figure S1A). These observations suggested that CL remodeling could respond to changes in mitochondrial lipids during long-term low oxygen exposure, a hallmark of natural yeast fermentation.

**Figure 1:**
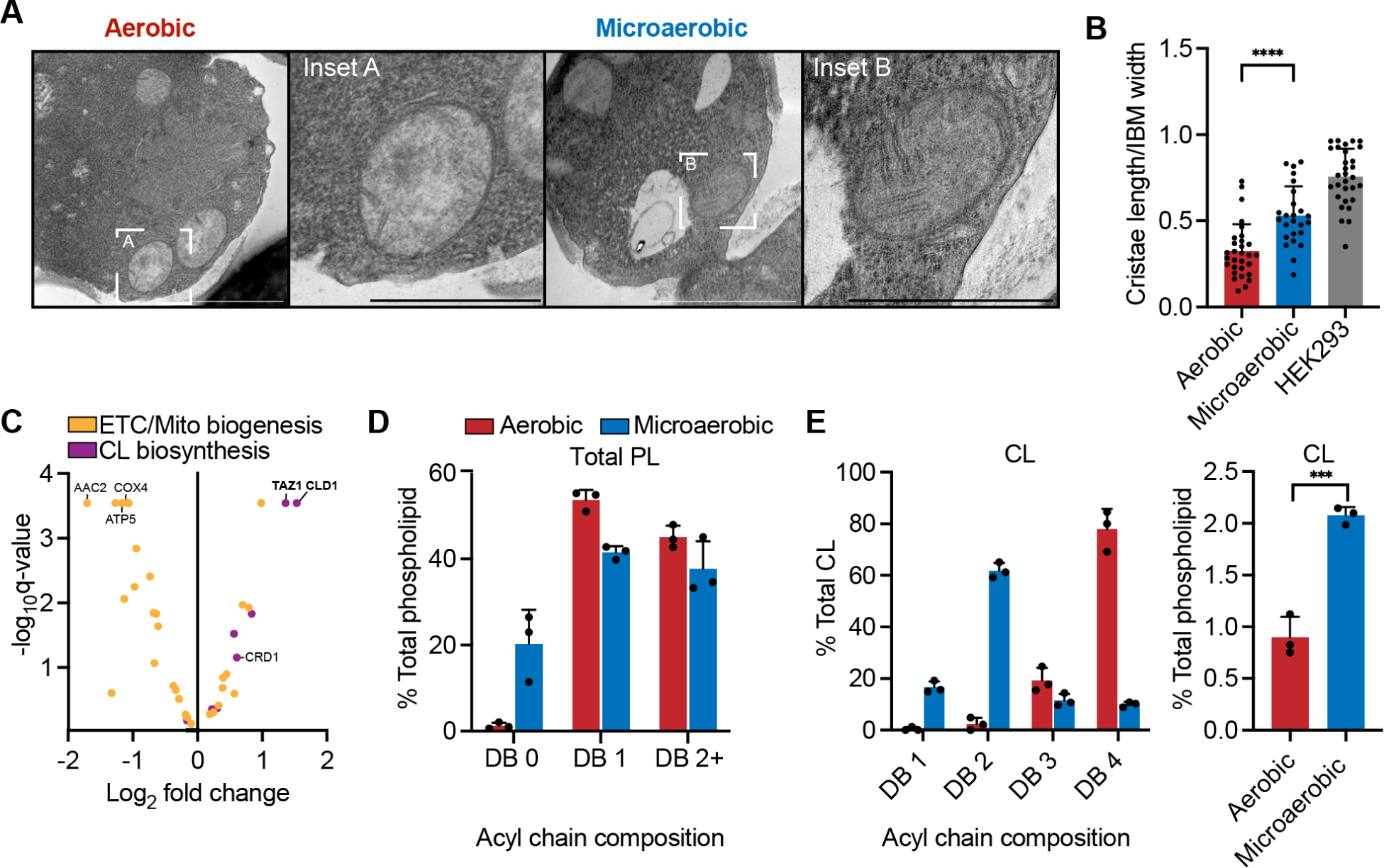
Microaerobic yeast feature elongated cristae structures and are enriched in CL. (**A**) Thin-section TEM images of mitochondria from yeast grown in glucose medium under aerobic and microaerobic conditions. Scale bars, 1000 nm. Insets scale bar, 500 nm. (**B**) Microaerobic cells have increased cristae length, more resembling those of the mammalian cell line HEK293. Cristae lengths were quantified relative to the width of the inner boundary membrane from N=30 mitochondria per condition. ****p<0.0001 unpaired t-test of aerobic vs. microaerobic yeast. Error bars indicate SD. (**C**) Gene expression changes in cells under microaerobic vs. aerobic growth implicates an increase in CL remodeling, but not CL biosynthesis. Reduced expression was observed for ETC components and proteins involved in mitochondrial biogenesis. RNA-seq data was originally collected and described in (66), data are available at the gene expression omnibus (accession number: GSE94345). (**D**) Microaerobic yeast cells possess more saturated PLs (DB 0) compared to aerobic cells as determined from lipidomic analysis on whole cells (n=3). Error bars indicate SD. (**E**) Left: Microaerobic yeast cells contain higher CL saturation (DB 1 and DB 2), but also an increase in total CL abundance (right). Relative levels of CL were determined by lipidomics (n=3). ***p=0.0007 unpaired t-test of aerobic vs. microaerobic cells.

Growth in low oxygen has been shown to increase the presence of saturated fatty acids through inhibition of fatty acid desaturases (67, 68). In yeast, a reduction in oxygen inhibits the activity of the sole lipid desaturase, Ole1p (55, 56). As a result, microaerobic yeast cells have reduced levels of unsaturated PLs (with one or two double bonds (DB) /PL) and increased levels of fully saturated PLs with 0 DB (Figure 1D). Elevated levels of saturated lipids have been associated with the induction of ER stress responses in mammalian cells, such as the unfolded protein response (UPR). However, we observed no elevated UPR activation or changes to ER morphology under microaerobic conditions (Figure S1B-C), suggesting that they do not represent a condition of fatty acid stress. Indeed, for highly abundant PLs like phosphatidylcholine (PC) or phosphatidylethanolamine (PE), microaerobic yeast lipidomes better mimic the acyl chain profile of mammalian cells with saturated *sn-1* and unsaturated *sn-2* chains (Figure S2), in contrast to di-unsaturated PLs in yeast grown under high aeration (54). In CL, we observed a major change in the unsaturation profile from predominantly 4 DB/CL in aerobic conditions to 2 DB in microaerobic conditions (Figure 1E). This change was coupled with a 2-fold increase in total CL abundance. Thus, microaerobic growth is characterized by a proliferation in mitochondrial cristae alongside an accumulation of saturated CL, potentially necessitating elevated activity of the remodeling pathway.

### Requirements for CL remodeling under microaerobic conditions driven by lipid saturation

We first asked if the increased levels of lipid saturation under microaerobic growth may provide an elevated role for CL remodeling in maintaining mitochondrial structure, as hinted by the increased expression of *CLD1* and *TAZ1* under these conditions (Figure 1C). In microaerobic conditions, we observed aberrant mitochondrial morphologies and reduced outgrowth in glucose for remodeling mutants, particularly in *cld1*Δ and *cld1*Δ*taz1*Δ, that were not present under aerobic growth (Figure 2A-C). TEM analysis also revealed shortened cristae tubules for *taz1*Δ and a complete absence of cristae structures in *cld1*Δ and *cld1*Δ*taz1*Δ cells in microaerobic conditions (Figure 2D). These cells instead featured flat IMMs that we previously observed in *crd1*Δ yeast, which lack CL, grown under microaerobic conditions or in backgrounds where lipid saturation was increased genetically (54). The effect of microaerobic growth on mitochondrial morphology was rescued by medium supplementation with oleic acid (Figure S3A-C), indicating that CL remodeling phenotypes under microaerobic conditions were the direct result of lipid saturation.

**Figure 2:**
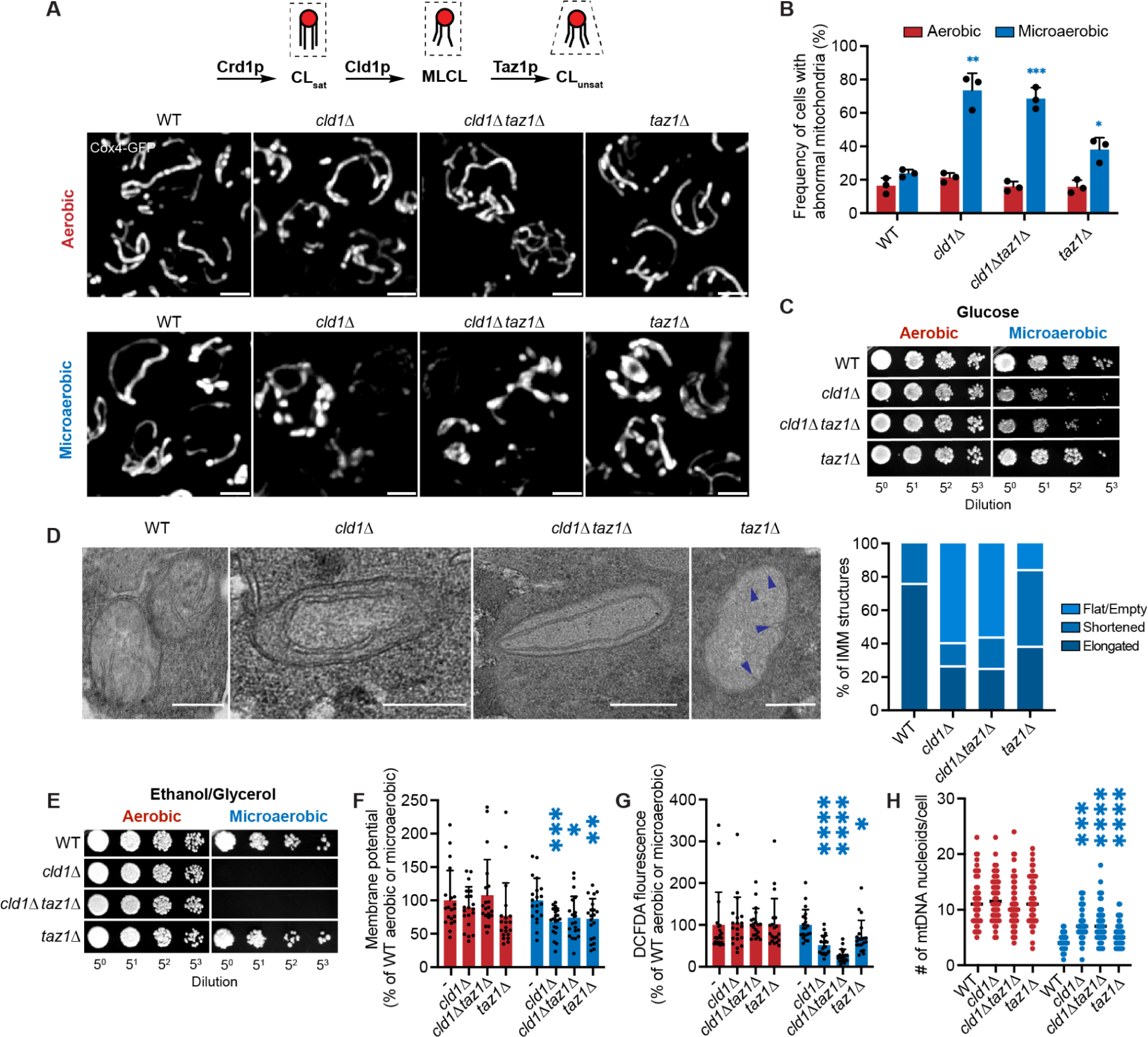
Loss of CL remodeling enzymes in microaerobic conditions results in altered IMM structure and mitochondrial dysfunction. (**A**) In aerobic conditions, loss of CL remodeling enzymes does not result in changes to mitochondrial morphology. However, in microaerobic conditions, *cld1*Δ*, cld1*Δ*taz1*Δ and *taz1*Δ cells show fragmented and swollen mitochondria. 3D projections of yeast cells expressing IMM-localized Cox4-GFP are shown. Scale bars, 2 μm. (**B**) Quantification of differences in the relative amounts of abnormal mitochondria in yeast strains bereft of CL remodeling enzymes grown in either aerobic or microaerobic conditions. N>50 cells were scored in each biological replicate. *p<0.05, **p<0.005, ***p<0.001 from unpaired t-tests against microaerobic WT and error bars indicate SD (n=3). (**C**) Loss of CL remodeling results in reduced fitness under microaerobic conditions. Liquid cultures were grown in aerobic (left) or microaerobic (right) conditions prior to serial dilution onto YPD agar plates, which were incubated under standard conditions. (**D**) CL remodeling mutants exhibit altered ultrastructure of cristae under microaerobic conditions. Left: thin section TEM of *cld1*Δ and *cld1*Δ*taz1*Δ microaerobic cells showed mitochondria with long, circular IMM structures that spanned the length of the OMM and resemble ‘flat’ cristae structures. Mitochondria from *taz1*Δ cells showed shortened cristae tubules (marked with arrows). WT cells possessed mitochondria with elongated (>200 nm) cristae sheets, as shown in Figure 1. Scale bars, 250 nm. Right: types of cristae/IMM observed by TEM were quantified from N>40 mitochondria in each condition. (**E**) Microaerobic cells lacking Cld1p are unable to grow on non-fermentable carbon sources. Liquid cultures in glucose medium were grown in either aerobic (left) or microaerobic conditions (right) prior to serial dilutions on YPEG agar plates, which were incubated under standard conditions. Only cells with functional mitochondria can form colonies on ethanol/glycerol as a carbon source in YPEG. (**F**) Mitochondrial potential was assayed under aerobic or microaerobic conditions. Maximum intensity of mitochondria in cells stained with 200 nM TMRE was measured using line profile analysis at the midplane of each cell, N=20 per condition. ***p=0.0008, *p=0.0162 and **p=0.0091 and unpaired t-test for *cld1*Δ, *cld1*Δ*taz1*Δ and *taz1*Δ respectively against WT. (**G**) Loss of CL remodeling dampens ROS levels under microaerobic conditions. ROS was assayed by microscopy after incubation of aerobic or microaerobic cells with 20μM H_2_DCFDA dye. Intensity was determined using line profile analysis at the midplane of each cell. N=20 cells were analyzed per condition. *p=0.0249, ****p<0.0001 unpaired t-test of *taz1*Δ, *cld1*Δ and *cld1*Δ*taz1*Δ respectively against WT. (**H**) Cells containing unremodeled CL exhibit increased numbers of mtDNA nucleoids compared to WT under microaerobic conditions. To image mtDNA nucleoids, cells were stained with SYBR Green I (SGI). The number of individual nucleoids in each cell were quantified using imageJ software, N=50 per condition. ***p=0.0005, ****p<0.0001 unpaired t-test of *cld1*Δ, *taz1*Δ and *cld1*Δ*taz1*Δ respectively against WT.

Significant defects in cristae architecture are often interlinked with respiratory dysfunction in the mitochondria. Since direct measurement of cellular respiration in microaerobic conditions is challenging, we instead tested other proxies of yeast mitochondrial function: outgrowth on non-fermentable carbon sources (ethanol/glycerol), membrane potential, reactive oxygen species (ROS) and mitochondrial nucleoid formation. While no changes to growth on ethanol/glycerol was observed in remodeling mutants in aerobic growth conditions, *cld1*Δ and *cld1*Δ*taz1*Δ were unable to grow after pre-culture in microaerobic conditions, indicating significant mitochondrial dysfunction (Figure 2E). Observed decreases in membrane potential, ROS, and an increase in mitochondrial nucleoids in all CL remodeling mutants also implied mitochondrial defects (Figure 2F-H). Akin to our examination of cristae structure, mitochondrial dysfunction was heightened in *cld1*Δ and *cld1*Δ*taz1*Δ backgrounds compared to *taz1*Δ, implying a greater role for loss of CL remodeling than accumulation of MLCL under microaerobic conditions.

### BTHS mutations drive mitochondrial abnormalities under microaerobic conditions

We next asked whether specific patient-derived BTHS mutants in Tafazzin could drive changes in mitochondrial morphology. While mitochondrial defects in *taz1*Δ were lower than those of *cld1*Δ and *cld1*Δ*taz1*Δ under microaerobic conditions, *taz1*Δ cells still exhibited significant perturbations compared to WT (Figure 2B, 2F-H). To test whether specific BTHS mutations accrued a similar effect on the mitochondria in these conditions, we utilized a previously described set of yeast strains that express *TAZ1* harboring mutations in conserved regions of human TAZ that also occur in BTHS (34, 35). Two BTHS-associated mutations, A88R and G230R, occur in the conserved acyltransferase domain and the membrane anchor region respectively. Importantly, A88R and G230R do not induce Taz1p mislocalization, but instead have defective Taz1p higher order structures and exhibit increased levels of MLCL and reductions in CL (34). Under aerobic conditions, BTHS mutants did not elicit alterations to mitochondrial morphology, function or ultrastructure (Figure 3A-D). However, under microaerobic growth, both mutants displayed altered IMM morphology, reduced potential and a loss of cristae ultrastructure (Figure 3A-D). These results demonstrate that BTHS-associated mutations can drive alterations to the yeast IMM when cells are grown under reduced oxygenation.

**Figure 3:**
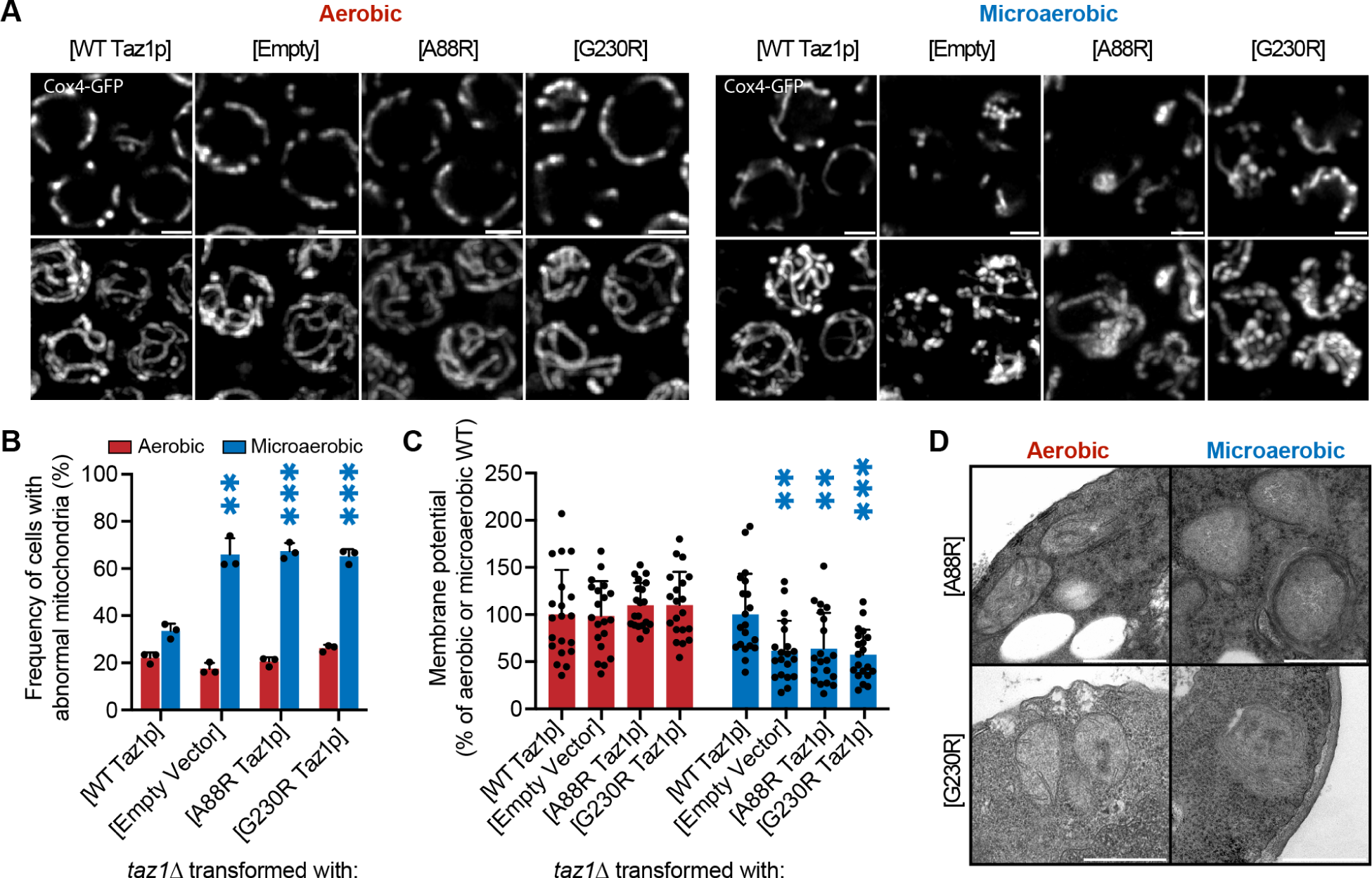
BTHS-associated Taz1p mutations exhibit abnormal mitochondrial architecture under microaerobic conditions. (**A**) Cells expressing BTHS mutations in conserved regions of Taz1p (BTHS mutants) show abnormal mitochondrial morphology under microaerobic, but not aerobic, growth conditions. Shown are example images of 2D and 3D projections of cells expressing IMM-localized Cox4-GFP. Scale bars, 2 μm. (**B**) Quantification of the relative amount of morphological defects in *taz1*Δ strains rescued with Taz1p harboring BTHS mutants compared to those rescued by WT Taz1p. N>50 cells were quantified in biological triplicates in either aerobic or microaerobic growth conditions. **p=0.0018 unpaired t-test of empty vector against WT in microaerobic conditions. ***p=0.0002 for A88R and G230R Taz1p against WT in microaerobic conditions. Error bars indicate SD. (**C**) Microaerobic growth results in reduced membrane potential in BTHS mutants compared to rescue. Cells were stained with 200 nM TMRE prior to imaging. ***p=0.0006 unpaired t-test of G230R Taz1p against WT Taz1p under microaerobic conditions. **p=0.0027 and 0.0078 unpaired t-test of WT against empty vector and A88R Taz1p respectively. (**D**) The BTHS rescue mutants, A88R and G230R, exhibit flat and empty inner membranes specifically under microaerobic conditions, while retaining cristae tubules under aerobic growth. Scale bars, 500 nm.

### Interactions between CL remodeling and lipid saturation are extendable to mammalian cells

With the strong interaction between Cld1p loss and microaerobic yeast growth, we asked whether inhibition of CL remodeling leads to loss of IMM ultrastructure in other cell types exposed to elevated lipid saturation. While no homolog of Cld1p exists in mammalian cells, the iPLA_2_ family of enzymes conducts an identical deacylation reaction on CL (21). We hypothesized that iPLA_2_ inhibition would elicit a similar sensitivity of mitochondrial structure to lipid saturation as we had observed in yeast. To test this, we treated HEK293 cells with the iPLA_2_ inhibitor bromoenol lactone (BEL), which has been shown previously to prevent MLCL formation by instead increasing the levels of nascent, unremodeled CL (22). To induce changes in lipid saturation, we fed cells with the saturated fatty acid Palm complexed to BSA at 100 μM, a concentration that does not induce any independent changes to mitochondrial functions (57, 69). We observed that BEL+Palm-treated cells exhibited respiratory dysfunction as well as perturbed mitochondrial morphology and ultrastructure (Figure 4A-C), which was not displayed in independent treatments of each compound. These results corroborate our findings in yeast and suggest that the inhibition of CL remodeling deacylation may be particularly sensitive to environmental saturated lipid levels.

**Figure 4:**
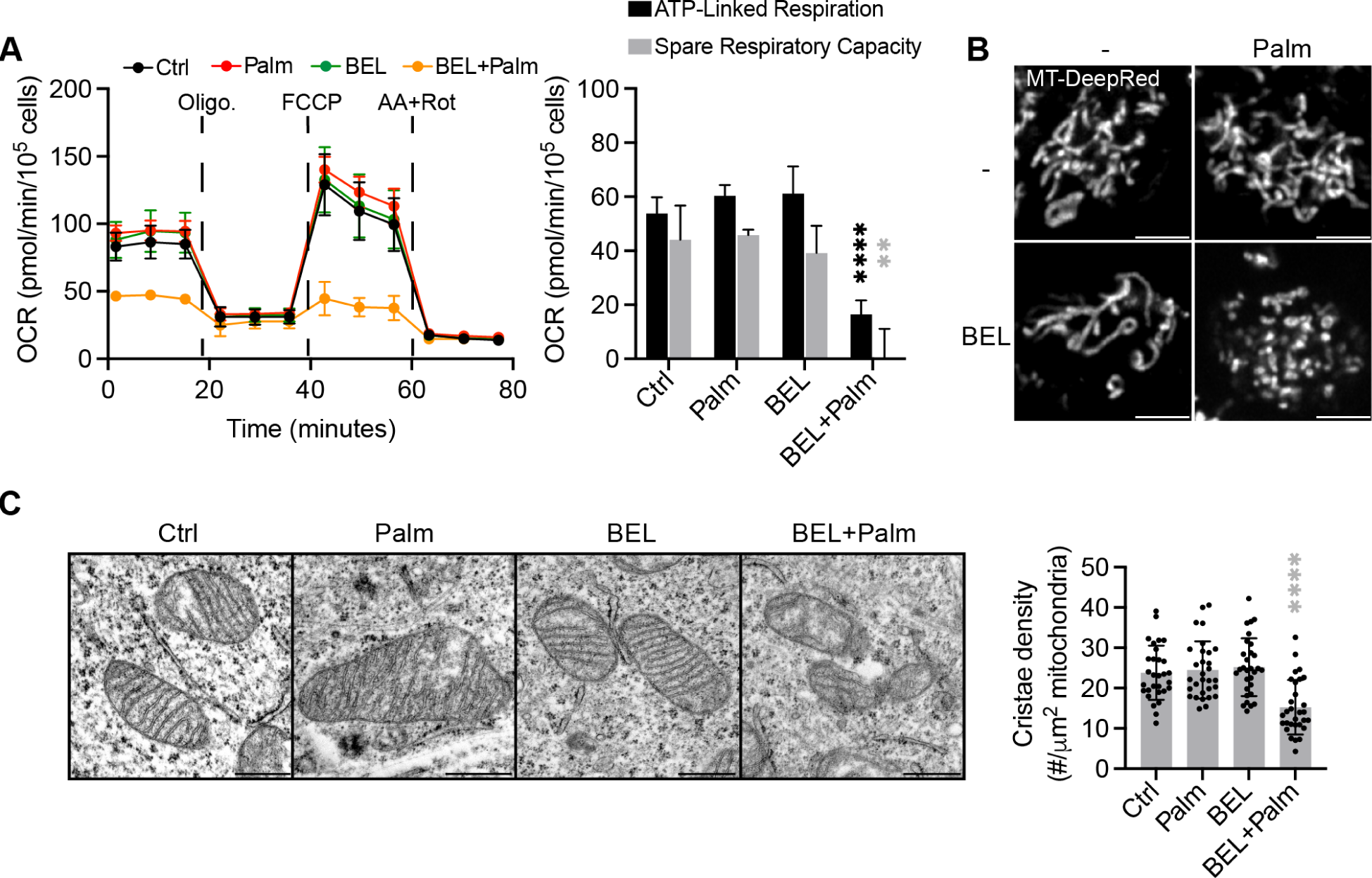
iPLA2 inhibition in saturated lipid environments results in mitochondrial dysfunction. (**A**) Seahorse respirometry of HEK293 cells treated with 2.5 μM bromoenol lactone (BEL) and 100 μM palmitate (Palm) results in reduced cellular respiration and a defective mitochondrial stress response. ATP-linked respiration is calculated from the decrease in OCR upon oligomycin injection. The spare respiratory capacity is an indicator of the cellular response to increased energetic demand and is defined as the difference between maximal and basal respiration. Defects in ATP-linked respiration and spare respiratory capacity are indicative of general ETC dysfunction in BEL+Palm cells. Wells containing 15000 cells were treated in biological replicates (n>3) for 48 hours prior to analysis. ****p<0.0001 and **p=0.0019 unpaired t-test of BEL+Palm against Ctrl for ATP-linked respiration and spare respiratory capacity respectively. Error bars indicate SD. (**B**) The combined treatment of BEL and Palm (BEL+Palm) results in fragmented mitochondria. HEK293 cells were treated as in (A) for 48 hours prior to imaging using 200 nM Mitotracker DeepRed FM. Scale bars, 5 μm. (**C**) Left: BEL+Palm results in empty mitochondria with few cristae, while all other conditions showed no defects in cristae density. Scale bars, 500 nm. Right: cristae density was measured for N=30 mitochondria in each condition. Error bars indicate SD. ****p<0.0001 unpaired t-test of BEL+Palm against Ctrl.

### Remodeling is coupled to total CL levels in microaerobic conditions

We next sought to delineate the lipidic drivers that led to mitochondrial dysfunction in yeast CL remodeling mutants under microaerobic, but not aerobic, growth conditions. Under aerobic conditions, we observed a ∼50% reduction in CL levels in each of the CL remodeling mutants compared to WT, and no changes to the CL precursor, phosphatidylglycerol (PG) (Figure 5A), in agreement with previous observations (47). Contrastingly, under microaerobic conditions, no CL was detected in *cld1*Δ or *cld1*Δ*taz1*Δ mutants and a ∼75% reduction in CL was observed in *taz1*Δ (Figure 5B). This loss in CL levels was coupled with increased PG, which suggested an inhibition of CL synthase. However, abrogation of CL synthesis in *crd1*Δ cells under microaerobic conditions produced much greater levels of PG, to ∼3% of all PLs (54), suggesting that inhibition of Crd1p is not the only factor contributing to loss of CL in *cld1*Δ or *cld1*Δ*taz1*Δ. We note that our analysis did not include quantification of MLCL, so we preclude that changes in its levels also occur under microaerobic growth.

**Figure 5:**
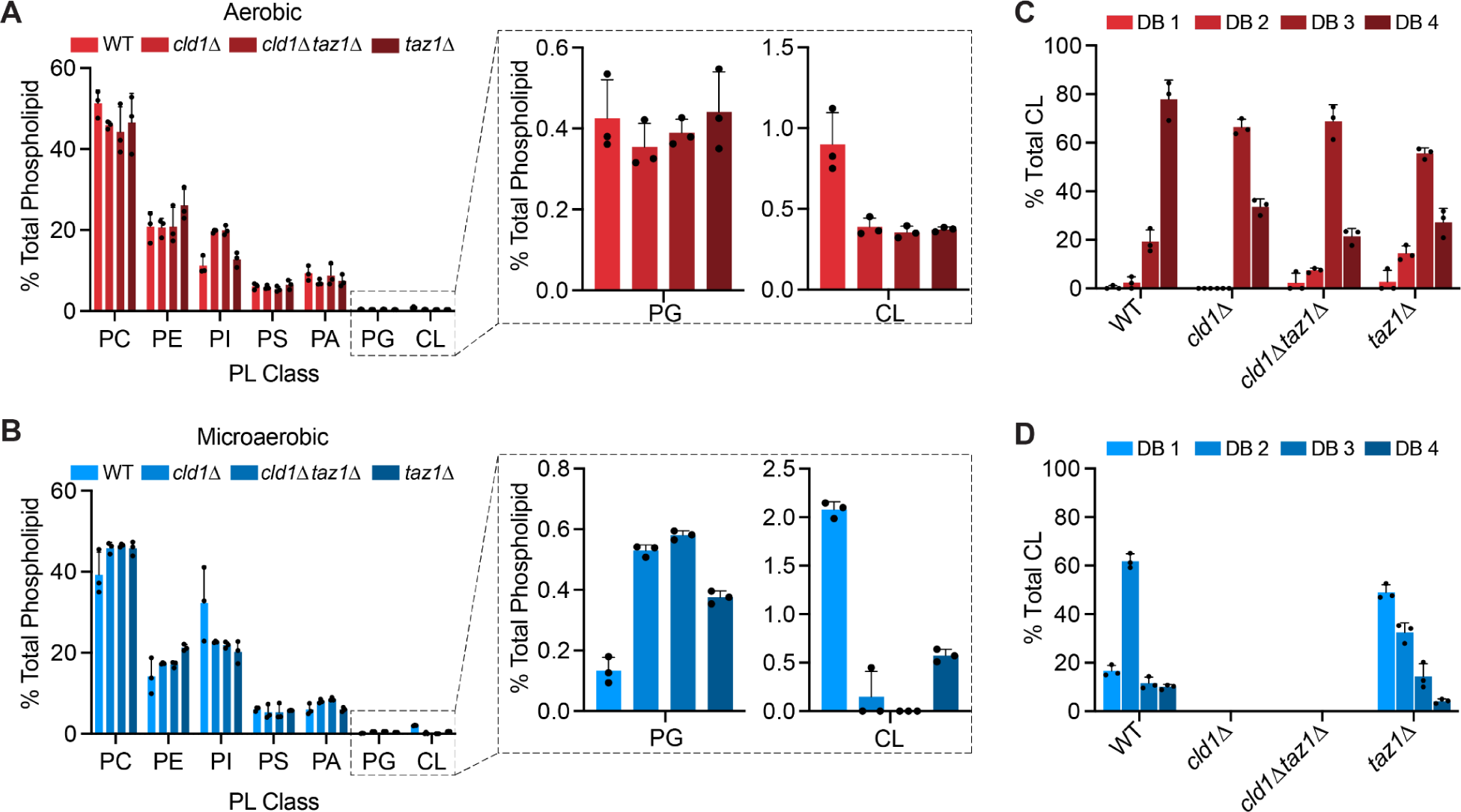
Differential lipidic changes under aerobic and microaerobic conditions in CL remodeling mutants. (**A**) Under aerobic conditions, removal of CL remodeling enzymes results in decreased CL levels without changes to the precursor PG. Error bars indicate SD (n=3). (**B**) Under microaerobic conditions, loss of CL remodeling enzymes results in a significant depletion of CL in *taz1*Δ cells and ablation of CL levels in *cld1*Δ and *cld1*Δ*taz1*Δ cells. A concomitant increase in PG levels was observed in each mutant, consistent with inhibition of CL synthase. Error bars indicate SD (n=3). (**C**) Aerobic CL remodeling mutants exhibit a shift in the CL double bond (DB) profile from 4 DB to 3 DB. Error bars indicate SD (n=3). (**D**) Microaerobic cells have more saturated CL molecules (4 DB to 2 DB), and removal of Taz1p shifts the saturation level of CL to predominantly 1 DB. No data is shown for *cld1*Δ and *cld1*Δ*taz1*Δ samples, as these predominantly had no detectable CL. Error bars indicate SD (n=3).

Further analysis of acyl chain composition revealed changes in CL resulting from the interplay of remodeling and saturation. Under aerobic conditions, *cld1*Δ and *cld1*Δ*taz1*Δ accumulate 3 DB CL instead of 4 DB CL; *taz1*Δ also accumulated 3 DB/CL as the principal CL species, but contained more saturated CL species with low levels of 2 or 1 DB (Figure 5C). Thus, all remodeling mutants under aerobic growth retain a highly unsaturated CL (3 DB) as the predominant species. In contrast, microaerobic growth altered the CL saturation levels from 4 DB/CL to 2 DB in WT cells, while the most abundant CL molecule in microaerobic *taz1*Δ was predominantly saturated (1 DB) (Figure 5D). These results aligned with our previous observations that high levels of genetically-induced lipid saturation induced significant mitochondrial defects, increased CL saturation to 1 DB/CL, and drove a 2-fold depletion in CL levels without changing to PG abundance or saturation (54). They suggest that a combination of increased CL saturation and reduced abundance could drive mitochondrial abnormalities under microaerobic conditions.

### Interactions between the phospholipase Ddl1p and the remodeling pathway

The loss of CL in *cld1*Δ backgrounds, but lack of accumulation of PG to *crd1*Δ levels, suggested that CL could be degraded when the remodeling pathway is absent under microaerobic conditions. We first considered whether mitophagic flux to the vacuole could drive CL degradation, as there is some evidence linking CL composition to mitophagy (70, 71). To test this, we measured the incorporation of mitochondrial Cox4-GFP intensity within the vacuole. As a comparison, we grew cells in glycerol for 72 hours, which induces mitophagy (72), and utilized a mitophagy-incapable *atg32*Δ background as a control for the inhibition of mitophagy (73). Under glycerol growth conditions, WT, *cld1*Δ, and *taz1*Δ all exhibited Cox4-GFP signal within the vacuole, while Cox4-GFP signal in *atg32*Δ was at the periphery of the cell and not in the vacuole (Figure S4). Neither WT nor CL remodeling mutants exhibited any vacuolar Cox4-GFP signal under microaerobic conditions. Thus, CL loss is unlikely to result from robust mitophagy under these conditions.

An alternative hypothesis is that CL can be specifically cleared through phospholipase activity. In BTHS patient-derived cell lines and *Drosophila*, defects in remodeling are proposed to drive enhanced turnover of CL via degradation from MLCL (36). However, loss of Cld1p, which prevents MLCL production, reduces CL levels in yeast, suggesting other mechanisms of breakdown. Apart from Cld1p, only the mitochondrial phospholipase A_1_, Ddl1p, has been implicated with putative CL hydrolase activity in yeast. Ddl1p has activity towards PA, PG, PC and PE as substrates (74, 75), but has also been shown to act on TOCL and MLCL (75).

We asked if knockouts of *DDL1* could restore mitochondrial function in remodeling mutants that accumulate reduced CL. Under aerobic conditions, *ddl1*Δ cells exhibited fragmented mitochondrial structures (Figure 6A), as has been previously described (75). Surprisingly, mitochondrial fragmentation was mitigated in *cld1*Δ*ddl1*Δ and *ddl1*Δ*taz1*Δ cells, suggesting an interaction between Ddl1p and the remodeling pathway. Under microaerobic conditions, *ddl1*Δ, *cld1*Δ*ddl1*Δ and *ddl1*Δ*taz1*Δ cells all exhibited significantly aberrant IMM morphologies (Figure 6A). However, short cristae tubules were observed in *cld1*Δ*ddl1*Δ under microaerobic conditions, unlike *cld1*Δ cells (Figure 6B). In microaerobic conditions, both *ddl1*Δ and *cld1*Δ*ddl1*Δ cells exhibited similar cristae structures and average lengths to *taz1*Δ (Figure 6B-C). When microaerobically-grown *cld1*Δ*ddl1*Δ cells were spotted on ethanol/glycerol plates, they regained viability that was lost in *cld1*Δ (Figure 6D). Loss of Ddl1p can therefore partially rescue IMM structure and function in *cld1*Δ backgrounds under microaerobic conditions.

**Figure 6:**
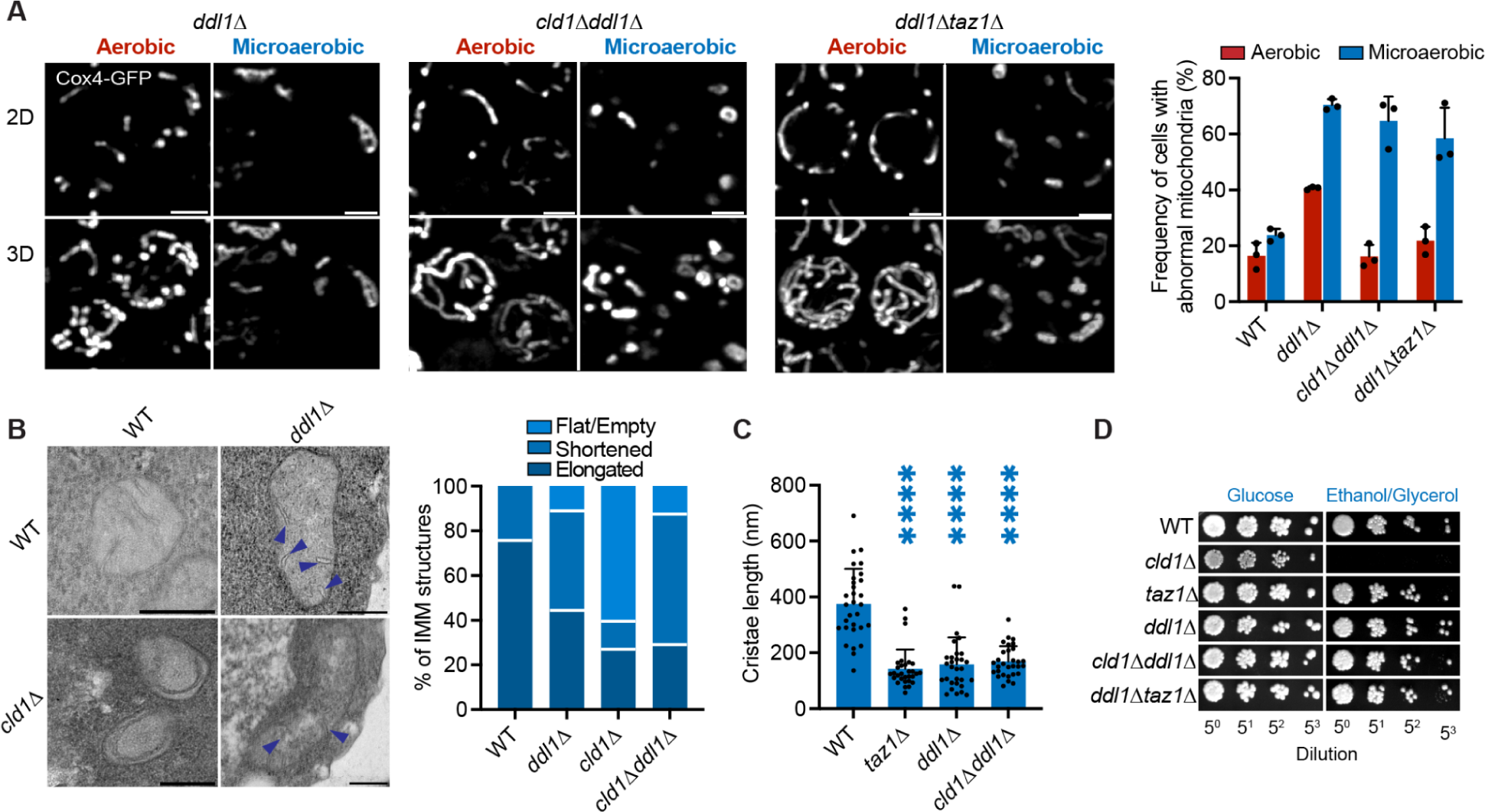
D*D*L1 deletion partially rescues cristae structure in *cld1*Δ backgrounds under microaerobic conditions. (**A**) Mitochondrial morphologies of *ddl1*Δ cells. Yeast expressing IMM-localized Cox4-GFP were grown in biological replicates (n=3) under aerobic or microaerobic conditions prior to imaging. Scale bars, 2 μm. For analysis of mitochondrial morphology (right), N>50 cells were quantified in each condition. All *ddl1*Δ strains exhibit elevated abnormal mitochondrial morphology under microaerobic growth, but removal of Cld1p or Taz1p reduces aerobic abnormality in the *ddl1*Δ background. **p=0.0096 and *p=0.0269 unpaired t-test for *cld1*Δ*ddl1*Δ and *ddl1*Δ*taz1*Δ against *ddl1*Δ respectively. Error bars indicate SD. (**B**) *ddl1*Δ cells have shortened cristae tubules as determined by thin-section TEM analysis. Arrows indicate shortened cristae tubules. Scale bars, 250 nm. In each condition N=40 mitochondria were quantified as containing either tubular, shortened or flat cristae. (**C**) *cld1*Δ*ddl1*Δ restores cristae tubules lost in microaerobic *cld1*Δ backgrounds, although they are of a shorter length than WT cells. Cristae length was quantified from N=30 mitochondria from each condition. ****p<0.0001 unpaired t-test of *taz1*Δ, *ddl1*Δ, *cld1*Δ*ddl1*Δ against WT. (**D**) Deletion of *DDL1* in *cld1*Δ backgrounds restores mitochondrial viability after microaerobic growth. Cells were grown in liquid medium under microaerobic conditions for 48 hours prior to serial dilution onto either YPD or YPEG plates, which were incubated at standard conditions.

We next conducted lipidomic analysis on aerobic and microaerobic cells of *ddl1*Δ in the presence and absence of CL remodeling mutants (Figure 7). We focused on the abundances and compositions of the PLs involved in the CL biosynthetic pathway: PA, PG and CL, all of which have been shown to be substrates for Ddl1p in vitro (Figure 7A). Under aerobic conditions, removal of *DDL1* in WT and CL remodeling-deficient backgrounds increased CL, but also PA and PG (Figure 7B). In *cld1*Δ and *taz1*Δ backgrounds, knockout of *DDL1* returned CL to WT levels, which may explain the rescue of mitochondrial morphology observed under aerobic growth (Figure 6A). Major increases in PA and PG were observed in *cld1*Δ*ddl1*Δ and *ddl1*Δ*taz1*Δ, but only minor changes in *ddl1*Δ (Figure 7B), suggesting that the increase in CL synthesis could be driven by increased PA in the absence of CL remodeling. Under microaerobic conditions, Ddl1p-deficient cells showed minimal CL levels in all backgrounds (Figure 7C). Notably, increased PG was observed in *ddl1*Δ and *taz1*Δ*ddl1*Δ, but not in *cld1*Δ*ddl1*Δ, suggesting increased flux towards CL synthesis in *cld1*Δ*ddl1*Δ. Increases in PA, however, were observed in all backgrounds (Figure 7C). While CL and PG are exclusively localized to mitochondria in yeast, PA is not, and we note that this analysis did not directly measure mitochondrial PA.

**Figure 7:**
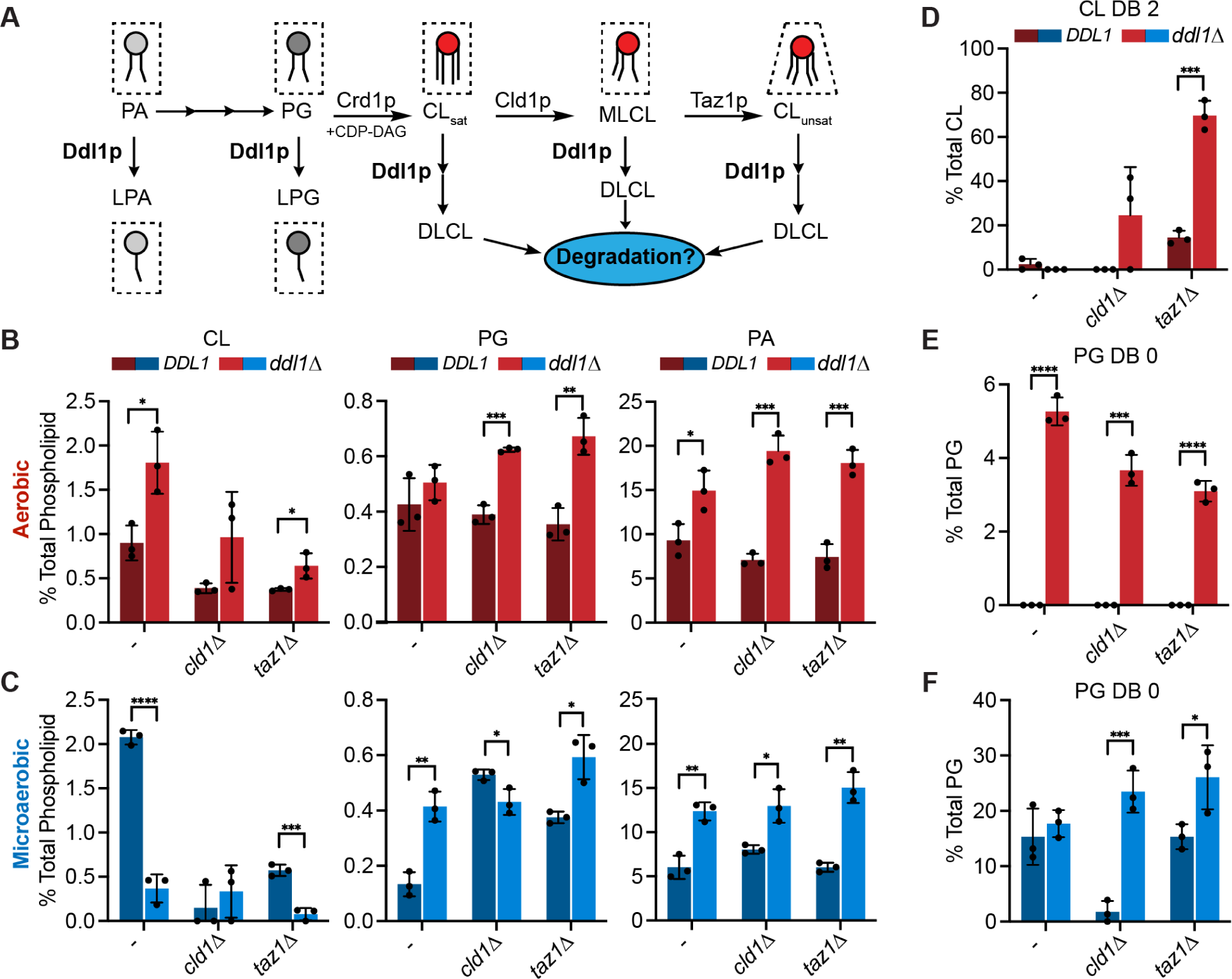
Ddl1p activity differentially affects PL metabolism in aerobic and microaerobic conditions. (**A**) Schematic depiction of the potential roles of phospholipase A_1_ activity in CL biosynthesis and remodeling pathways based on previously observed biochemical activities of Ddl1p or its mammalian homologue. (**B**) Loss of *DDL1* in CL remodeling-deficient backgrounds results in increased PA and PG in aerobic conditions, and a smaller increase in CL levels. For all subsequent analyses in this figure: Error bars indicate SD (n=3) and *p<0.05, **p<0.005, ***p<0.001 and ****p<0.0001 from unpaired t-tests from each background ± *DDL1*. (**C**) Under microaerobic conditions, *ddl1*Δ backgrounds exhibit reduced CL levels, but maintain increases in PG and PA. (**D**) Saturated CL (DB 2) appears in remodeling mutants lacking Ddl1p grown under aerobic conditions. (**E**) Fully saturated PG (DB 0) appears in WT and remodeling mutants lacking Ddl1p grown under aerobic conditions. (**F**) Fully saturated PG (DB 0) increases only in remodeling mutants lacking Ddl1p grown under microaerobic conditions.

In addition to modifying PL biosynthesis, loss of Ddl1p drove differential changes to PL acyl chain composition under aerobic and microaerobic growth. In aerobic conditions, *DDL1* deletion affected the CL chain composition of remodeling mutants. Most notably, a large increase in DB 2 CL was observed in *ddl1*Δ*taz1*Δ compared to *taz1*Δ cells (Figure 7D). There was also a consistent appearance of fully saturated PG in each of the *ddl1*Δ mutants, which were not observed in any of the parental strains (Figure 7E). Under microaerobic conditions, PG and PA saturation levels also increase in *ddl1*Δ backgrounds (Figure 7F, Figure S5B). Minor changes were observed to CL saturation under microaerobic conditions, but the low overall levels of CL made these difficult to define (Figure S5B). Taken together, these data suggest that Ddl1p could modulate CL levels through activity on its precursors PA and PG.

## Discussion

In this study, we address why removal of CL remodeling enzymes do not render any morphological or ultrastructural phenotype in yeast (47). We showed that CL remodeling is essential for mitochondrial structure and function under microaerobic growth conditions, in which the yeast lipidome is defined by increases in lipid saturation. Microaerobically-grown remodeling mutants exhibited a loss of highly curved cristae in the IMM, which resemble structures observed in BTHS patients (32) but are not seen in aerobically-grown yeast mutants. Phenotypes were especially strong in *cld1*Δ cells that show a depletion in CL levels. We extended this observation to mammalian cells, where the abundance of cristae in BEL-treated HEK293 was reduced upon a mild treatment of exogenous saturated fatty acids. These data support a model that the role of CL remodeling can be dependent on the overall fatty acid metabolism of cells. They also suggest that the function of the yeast remodeling pathway could be to act during microaerobic growth, a natural condition that promotes lipid saturation.

Yeast cells grown under vigorous laboratory aeration have high levels of unsaturated lipids, which may mask the defects associated with the removal of CL remodeling enzymes. Under aerobic conditions, remodeling mutants retain high levels of CL unsaturation (3 DB/CL) because the substrates for CL synthesis are predominantly di-unsaturated (Figure S2). Under microaerobic conditions, CL in *taz1*Δ is predominantly saturated (1 DB) and CL is lost in *cld1*Δ (Figure 5B, 5D). These differences could have significant effects on the properties of CL. While fully unsaturated CL (TOCL) can adopt high-curvature non-lamellar phases (76–78), fully saturated – tetra-palmitic (16:0) or tetra-myristic (14:0) – CL exhibit gel phases (78, 79) that are likely unsuitable for functions in the IMM. The number of saturated acyl chains incorporated in CL could thus drive significant changes in its biophysical properties, which may be crucial for proper cristae assembly. Proteins like MICOS and ATP synthase also contribute to IMM curvature (80–82) and their decreased expression during microaerobic conditions might act as an additional factor in increasing the dependence on lipid-encoded curvature for shaping IMM morphology. The predominant unsaturated form of MLCL (47) also adopts lamellar phases, indicating a reduced curvature than remodeled CL (83–85), and fails to interact with ETC complexes and supercomplexes.

A general paradigm of BTHS pathophysiology is that MLCL levels are the primary drivers of ETC dysfunction and reduced respiratory capacity observed in patients (86, 87). However, mitigation of the MLCL:CL ratio through BEL treatment still results in perturbed respiratory capacity in several BTHS models (21, 39, 40). BEL-treated cells have been shown to have large amounts of unremodeled CL (22), similar to *cld1*Δ yeast. Our results with *cld1*Δ cells under microaerobic conditions and the combined effects of Palm and BEL-treatment in HEK 293 cells support a hypothesis that the accumulation of unremodeled (saturated) CL could act as an additional contributor to mitochondrial dysfunction. If so, availability of exogenous fatty acids might be a key regulator of phenotypes in CL remodeling mutants. Indeed, previous studies in HeLa cell knockouts of TAZ have observed a sensitivity of CL and mitochondrial function to changing saturated fatty acid levels (88, 89). Conversely, linoleic acid (18:2) treatment of a BTHS iPSC cardiomyocyte model provided a comparatively greater rescue of mitochondrial functionality than BEL (40), but was less effective in the restoration of cardiac function in a BTHS mouse model (90). Further analyses across a range of systems would be needed to disentangle the individual contributions of MLCL and unremodeled CL to mitochondrial dysfunction. Notably, the analyses performed in this study do not quantify MLCL, and therefore we cannot preclude its role in phenotypes reported here.

The interplay of CL remodeling, biosynthesis, and potential degradation pathways are another area of future work. We observed that loss of the mitochondrial phospholipase A_1_ Ddl1p partially rescued cristae structures in *cld1*Δ backgrounds under microaerobic growth (Figure 6). We originally hypothesized that Ddl1p could act to degrade unremodeled CL, leading to the loss of CL in microaerobic *cld1*Δ cells. While *ddl1*Δ cells showed increased CL levels in aerobic growth (Figure 7B), they did not restore CL levels under microaerobic conditions (Figure 7C). Instead, increased abundance and saturation of PA and PG, which are precursors to CL, were observed in all conditions. One possibility is that the accumulation of saturated PG and PA may prevent CL synthesis under microaerobic conditions, as they are poor substrates for CL synthase and CDP-DAG synthase, respectively (91). We previously observed that high levels of genetically-induced lipid saturation increased CL saturation to 1 DB/CL, and drove a 2-fold decrease in its abundance, albeit without an accumulation of PG or PA (54). Interestingly, both CL and PL saturation control the formation of non-lamellar lipid topologies, which are required for optimal TAZ activity (23), but could also be relevant for the activity of other steps in CL metabolism.

An open question remains why loss of Ddl1p rescues mitochondrial viability under microaerobic conditions. One possibility is that increased levels of the anionic precursors PA or PG might fulfill some of the structural roles of CL. Testing this model would be aided by biochemical analyses of purified mitochondria from microaerobic cells, which currently remains a challenge. Further experiments are also required to better define the phospholipase activity of Ddl1p in cells. The mammalian homologs of Ddl1p, DDHD1/2, have been described as preferentially hydrolyzing PA (92), but activities towards CL have not been reported. Mutations in DDHD1 cause a subtype of hereditary spastic paraplegia (HSP), which specifically exhibits defects in respiratory capacity and increases in oxidative stress (93), raising the possibility that DDHD1 may also alter CL composition, either directly or indirectly through action on CL substrates. In yeast, Ddl1p might also be involved in the heightened levels of CL we observe under microaerobic conditions. These were surprising given the established correlation between CL synthesis and OXPHOS components (19) and the lack of change in mitochondrial volume we have previously measured under identical conditions (54). Aerobic *ddl1*Δ cells accumulate CL to similar levels (∼2% of a PLs in the cell) as microaerobic WT cells, suggesting that Ddl1p might be responsible for regulating CL levels in response to oxygenation. Previous work has demonstrated that Aft1p, a transcriptional activator that is activated in response to low intracellular iron associated with hypoxia (94), negatively regulates *DDL1* expression and its deletion causes decreased levels of CL (75). Microaerobic growth could thus lead to activation of Aft1p, repressing *DDL1* expression and increasing CL levels.

Our data supports a model that while CL remodeling is dispensable for IMM structure and function in very unsaturated lipid environments, it is likely required to maintain the structure of cristae under conditions of elevated lipid saturation. Both this study and our previous work (54) have demonstrated the sensitivity of CL synthesis and remodeling to changes in the lipidic environment, as both processes become essential for yeast mitochondria under conditions of increased saturation, which has also been alluded to in mammalian cells (89). Such a dynamic could be relevant for understanding the cell-type specific phenotypes of BTHS that arise despite the loss of CL remodeling across a wide range of tissues (95). It could also play a role in the heterogeneity in BTHS pathophysiology observed in populations harboring TAZ mutations (95), given the myriad of dietary and metabolic factors that control the composition of circulating and cellular fatty acids. The phenotypes of the remodeling pathway mutants reported here could thus inform the interactions between CL and fatty acid metabolism contributing to mitochondrial dysfunction.

## Data Availability

All data are contained within the article and the supplemental data. All strains and plasmids are available upon request to the corresponding author.

## Supplemental data

This article contains supplemental figures S1-5 and a spreadsheet of lipidomics data.

## Abbreviations

IMM: Inner mitochondrial membrane
PL: Phospholipid
ETC: Electron Transport Chain
CM: Cristae membrane
CL: Cardiolipin
MLCL: Monolysocardiolipin
PE: Phosphatidylethanolamine
PC: Phosphatidylcholine
PI: Phosphatidylinositol
PG: Phosphatidylglycerol
PS: Phosphatidylserine
PA: Phosphatidic acid
BTHS: Barth syndrome
Palm: Palmitic Acid
TAZ: Tafazzin
OA: Oleic acid
BEL: Bromoenol lactone
iPLA_2_: Intracellular Phospholipase A_2_.

## Acknowledgements

The authors would like to thank Steven Claypool, Vishal Gohil, Miriam Greenberg and Budin lab members for their helpful discussions. Steven Claypool generously provided yeast strains. Sterling Ramsey assisted in initial strain development. The National Institutes of Health (NIH) (R35-GM142960 to IB) and the Department of Energy (DE-SC0022954 to IB) provided financial support. KV was supported by the NIH Molecular Biophysics Training Grant (T32-GM008326C). The UCSD-CMM-EM Core (RRID: SCR_022039), partly supported by NIH S10OD023527, provided technical assistance and equipment access.

## Supplementary Information

**Figure S1:**
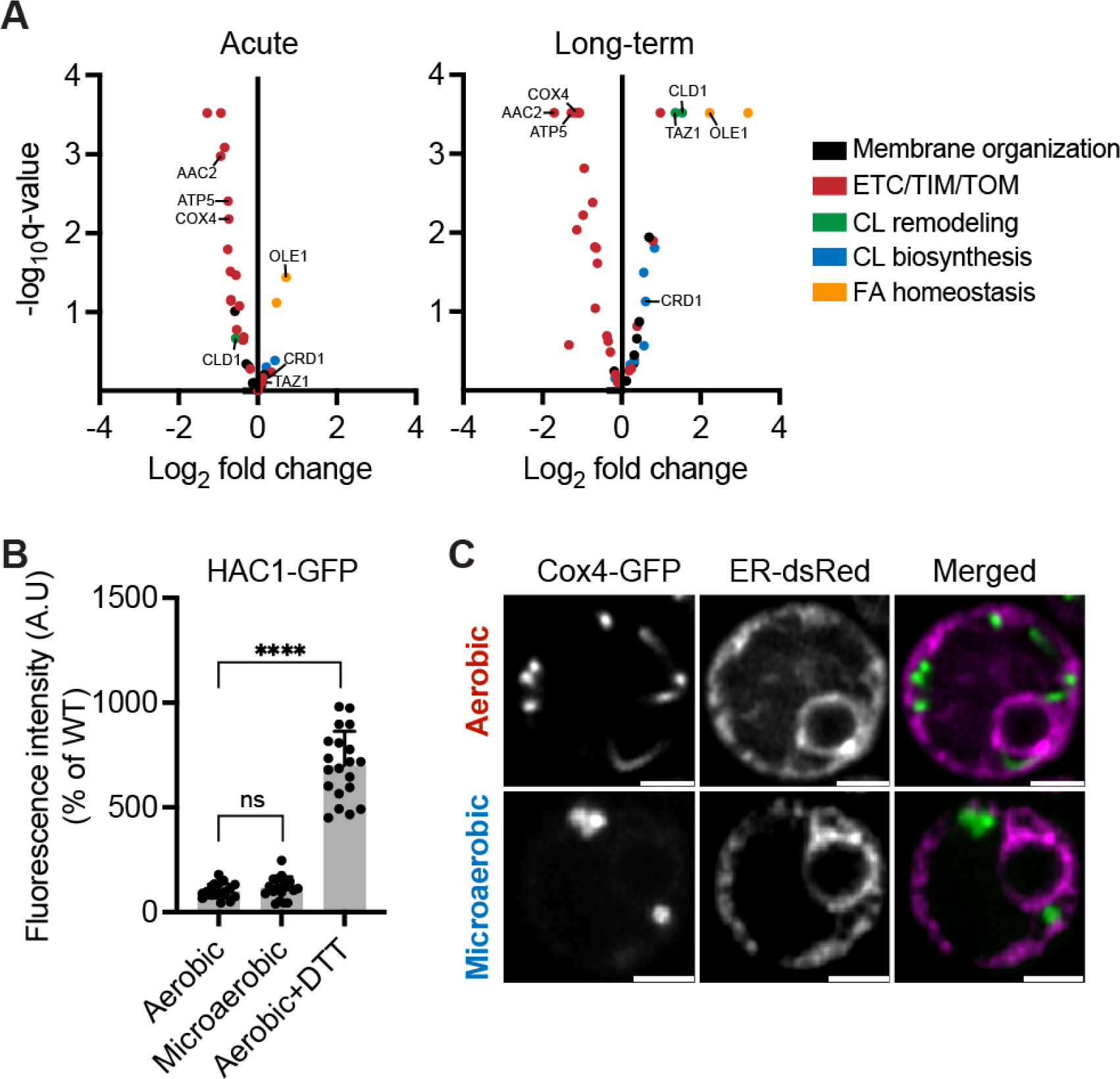
Microaerobic growth upregulates CL remodeling but does not induce ER stress. (**A**) Volcano plots showing gene expression changes between aerobic and microaerobic growth conditions upon acute (short-term) and extended (2 days) oxygen deprivation. RNA-seq data was originally collected and described in (66), data are available at the gene expression omnibus (accession number: GSE94345). (**B**) Microaerobic growth conditions do not induce UPR activation. ER stress was measured using a HAC1-GFP splice reporter. GFP intensity was measured from N=20 cells. WT cells treated with 5 mM dithiothreitol (DTT) were used as a control that elicits a strong UPR response. ****p<0.0001 from an unpaired t-test of aerobic+DTT against the untreated control. Error bars indicate SD. (**C**) Microaerobic growth does not elicit morphological changes to ER structure in cells expressing dsRed bearing a KDEL localization tag (ER-dsRed). Scale bars, 2 μm.

**Figure S2:**
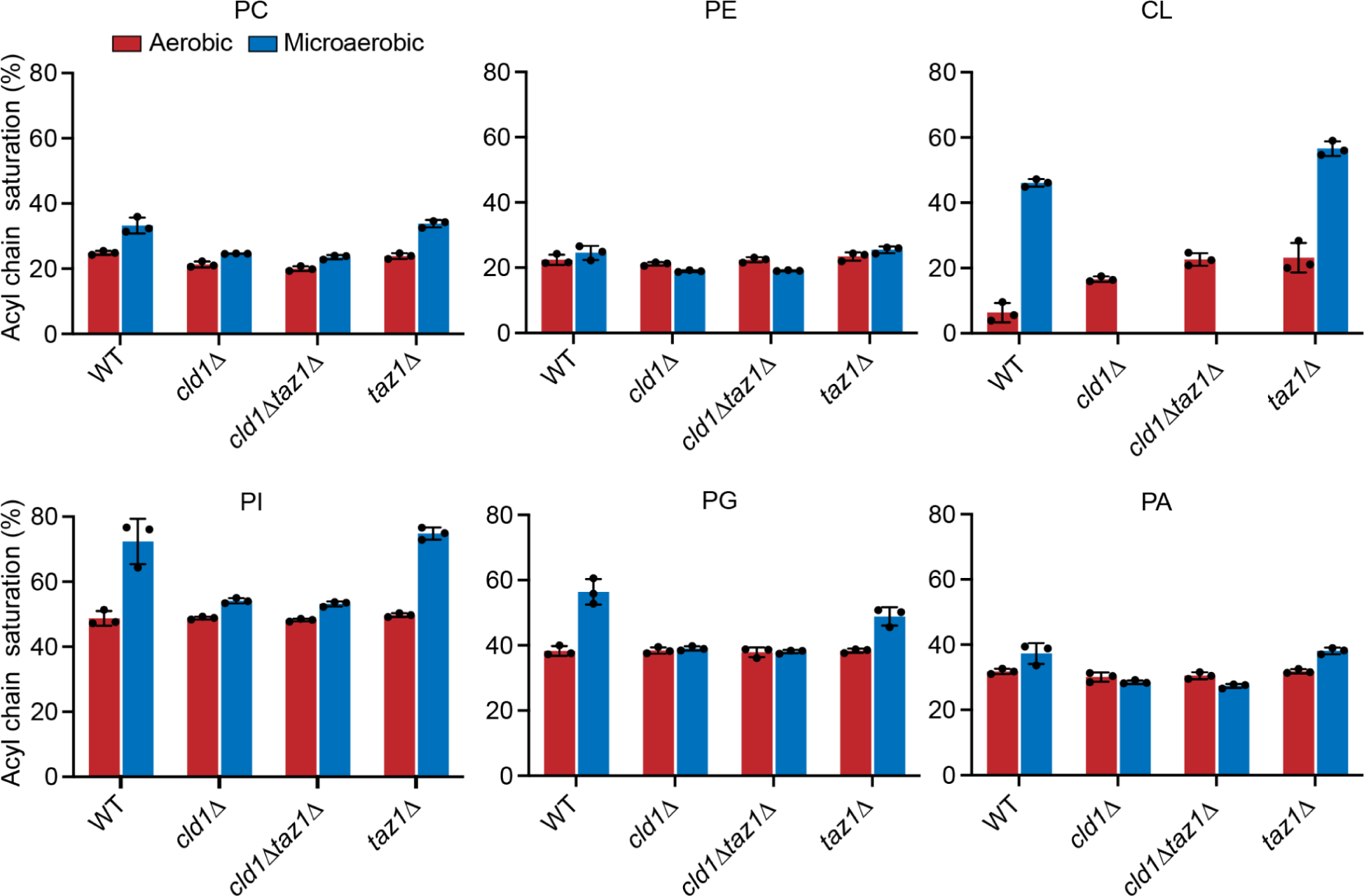
Microaerobic growth results in changes to the incorporation of saturated fatty acids (SFAs) species into multiple PL classes, but CL shows the largest changes. The acyl chain saturation % refers to the relative incidence of a PL acyl chain being saturated (0 DB), i.e an acyl chain saturation of 50% means half of the acyl chain pool does not contain a double bond. Lipidomics analysis was conducted in biological triplicates (n=3) grown in either aerobic or microaerobic conditions. Error bars indicate SD.

**Figure S3:**
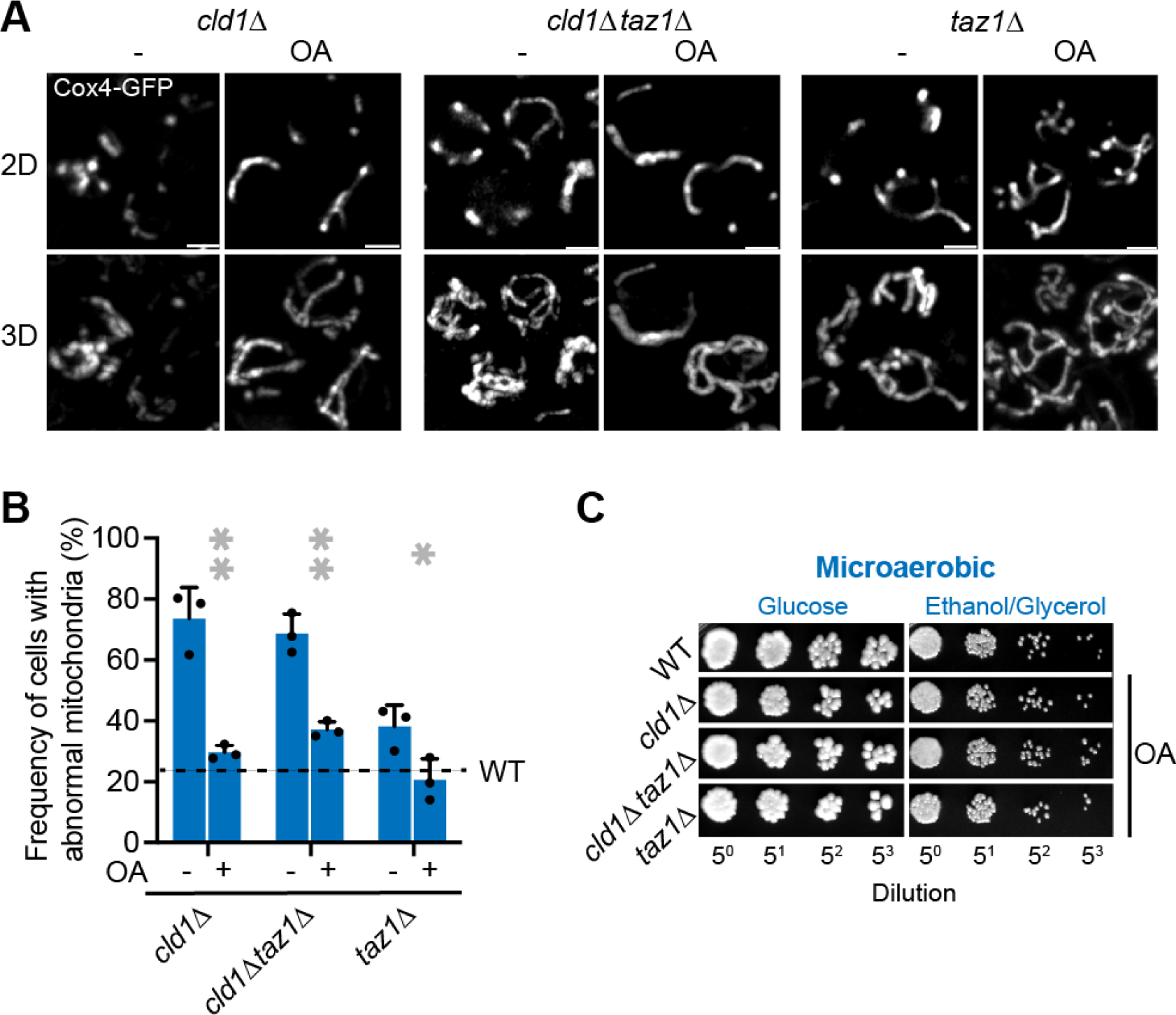
CL remodeling specifically interacts with increasing lipid saturation. (**A**) & (**B**) Oleic acid (OA) treatment of CL remodeling mutants under microaerobic conditions rescues aberrancies in mitochondrial morphology. **p=0.0026 and 0.0047 unpaired t-tests for OA-treated *cld1*Δ and *cld1*Δ*taz1*Δ against their untreated counterparts. *p=0.0368 unpaired t-test of OA-treated *taz1*Δ against the untreated condition. Error bars indicate SD. Scale bars, 2 μm. (**C**) OA-treatment restores *cld1*Δ and *cld1*Δ*taz1*Δ growth on ethanol/glycerol to WT levels after microaerobic growth. Cells were grown in biological triplicates (n=3) in medium supplemented with 356 μM OA solubilized in tergitol. Growth occurred in liquid culture for 2 days in microaerobic conditions followed by serial dilution on either glucose or ethanol/glycerol plates, which were then incubated under standard conditions.

**Figure S4:**
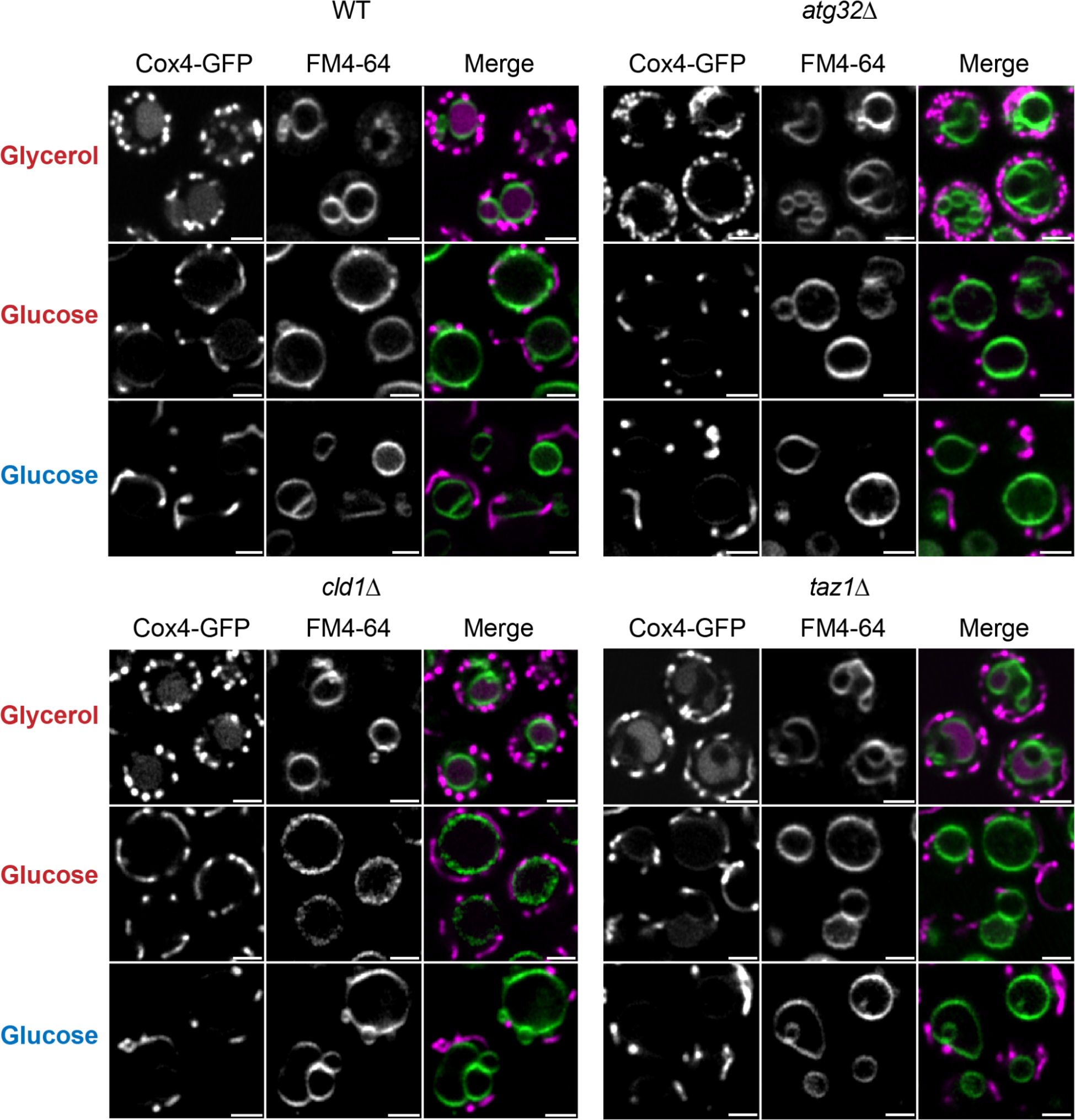
Loss of CL remodeling under microaerobic conditions does not induce mitophagy. Under conditions that trigger mitophagic flux to the vacuole, i.e glycerol growth for 72 hours, mitochondrial Cox4-GFP signal is observed in the vacuole. In *atg32*Δ conditions, mitophagy is inhibited, thus preventing any vacuolar Cox4-GFP signal. Microaerobic growth, like aerobic glucose growth, did not trigger mitophagy, as evidenced by a lack of Cox4-GFP signal in the vacuole. No changes between WT and CL remodeling strains was observed under microaerobic conditions, implying an alternative mechanism for depletion of CL. Scale bars, 2 μm.

**Figure S5:**
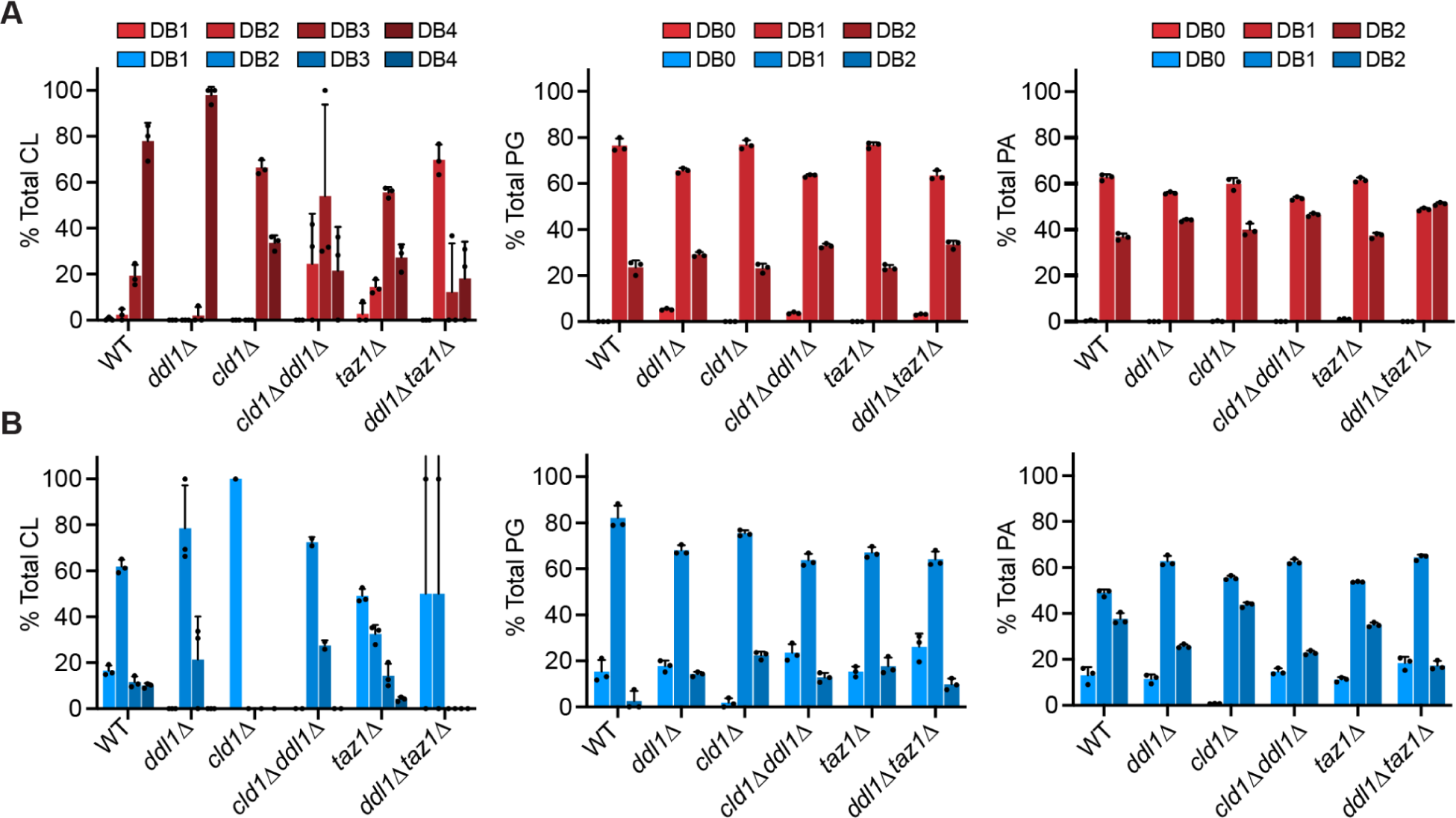
Changes to anionic PL saturation in response to loss of *DDL1* in CL remodeling mutants. (**A**) Lipidomic analysis of CL, PG and PA acyl chain compositions under aerobic conditions reveals that loss of *DDL1* specifically increases saturated CL species (DB1 and DB 2) in CL remodeling-deficient backgrounds (*cld1*Δ*ddl1*Δ and *ddl1*Δ*taz1*Δ). Error bars indicate SD (n=3). (**B**) Under microaerobic conditions, *DDL1* deletion increases saturated PG and PA (DB 0) in *cld1*Δ backgrounds. Error bars indicate SD (n=3).

**Table S1:** Yeast (*S. cerevisiae*) strains used in this study. Experimental Models: Organisms/strains

Experimental Models: Organisms/strains
| Reagent or Resource | Source | Description |
| --- | --- | --- |
| W303a (*MATa leu2-3112 trp1-1 can1-100 ura3-1 ade2-1 his3-11,15*) | ATCC | Haploid background strain |
| W303a, *taz1Δ::kanMX4* | This study |  |
| W303a, *cld1Δ::trp1* | This study |  |
| W303a, *cld1Δtaz1Δ- cld1Δ::trp1, taz1Δ::kanMX4* | This study |  |
| W303a, *atg32Δ::his3MX6* | This study |  |
| W303a, *ddl1Δ::kanMX4* | This study |  |
| W303a, *cld1Δ::trp1, ddl1Δ::kanMX4* | This study |  |
| W303a, *ddl1Δ::kanMX4, taz1Δ::his3MX6* | This study |  |
| GA-74-1A (*MATa leu2, ura3, trp1, ade8*)<br>*taz1Δ::HISMX6* | (34, 35) | Parent strain for BTHS mutant panel. Transformed with either WT Taz1, Empty vector, A88R Taz1 and G230R Taz1. |

**Table S2:** Plasmids used in this study.

| Reagent or Resource | Source | Description |
| --- | --- | --- |
| Cox4-GFP | Dr. Zhiping Xie | CIV-subunit 4-GFP, IMM-localized |
| pPW599 | Dr. Peter Walter | UPR splice reporter |
| pPW1409 | Dr. Peter Walter | ER-localized RFP |

## References

1. Acoba, M. G., N. Senoo, and S. M. Claypool. 2020. Phospholipid ebb and flow makes mitochondria go. J. Cell Biol. 219. [online] 10.1083/jcb.202003131.

2. Pfeiffer, K., V. Gohil, R. A. Stuart, C. Hunte, U. Brandt, M. L. Greenberg, and H. Schägger. 2003. Cardiolipin stabilizes respiratory chain supercomplexes. J. Biol. Chem. 278: 52873–52880.

3. Gohil, V. M., P. Hayes, S. Matsuyama, H. Schägger, M. Schlame, and M. L. Greenberg. 2004. Cardiolipin biosynthesis and mitochondrial respiratory chain function are interdependent. J. Biol. Chem. 279: 42612–42618.

4. Ren, M., C. K. L. Phoon, and M. Schlame. 2014. Metabolism and function of mitochondrial cardiolipin. Progress in Lipid Research. 55: 1–16. [online] 10.1016/j.plipres.2014.04.001.

5. Paradies, G., V. Paradies, V. De Benedictis, F. M. Ruggiero, and G. Petrosillo. 2014. Functional role of cardiolipin in mitochondrial bioenergetics. Biochimica et Biophysica Acta (BBA) – Bioenergetics. 1837: 408–417. [online] 10.1016/j.bbabio.2013.10.006.

6. Beltrán-Heredia, E., F.-C. Tsai, S. Salinas-Almaguer, F. J. Cao, P. Bassereau, and F. Monroy. 2019. Membrane curvature induces cardiolipin sorting. Communications Biology. 2. [online] 10.1038/s42003-019-0471-x.

7. Ikon, N., and R. O. Ryan. 2017. Cardiolipin and mitochondrial cristae organization. Biochimica et Biophysica Acta (BBA) – Biomembranes. 1859: 1156–1163. [online] 10.1016/j.bbamem.2017.03.013.

8. DeVay, R. M., L. Dominguez-Ramirez, L. L. Lackner, S. Hoppins, H. Stahlberg, and J. Nunnari. 2009. Coassembly of Mgm1 isoforms requires cardiolipin and mediates mitochondrial inner membrane fusion. J. Cell Biol. 186: 793–803.

9. Bustillo-Zabalbeitia, I., S. Montessuit, E. Raemy, G. Basañez, O. Terrones, and J.-C. Martinou. 2014. Specific interaction with cardiolipin triggers functional activation of Dynamin-Related Protein 1. PLoS One. 9: e102738.

10. Stepanyants, N., P. J. Macdonald, C. A. Francy, J. A. Mears, X. Qi, and R. Ramachandran. 2015. Cardiolipin’s propensity for phase transition and its reorganization by dynamin-related protein 1 form a basis for mitochondrial membrane fission. Mol. Biol. Cell. 26: 3104–3116.

11. Dudek, J.. 2017. Role of Cardiolipin in Mitochondrial Signaling Pathways. Front Cell Dev Biol. 5: 90.

12. Paradies, G., V. Paradies, F. M. Ruggiero, and G. Petrosillo. 2019. Role of Cardiolipin in Mitochondrial Function and Dynamics in Health and Disease: Molecular and Pharmacological Aspects. Cells. 8: 728. [online] 10.3390/cells8070728.

13. Hoch, F. L.. 1992. Cardiolipins and biomembrane function. Biochim. Biophys. Acta. 1113: 71–133.

14. Schlame, M., M. Ren, Y. Xu, M. L. Greenberg, and I. Haller. 2005. Molecular symmetry in mitochondrial cardiolipins. Chem. Phys. Lipids. 138: 38–49.

15. Oemer, G., J. Koch, Y. Wohlfarter, M. T. Alam, K. Lackner, S. Sailer, L. Neumann, H. H. Lindner, K. Watschinger, M. Haltmeier, E. R. Werner, J. Zschocke, and M. A. Keller. 2020. Phospholipid Acyl Chain Diversity Controls the Tissue-Specific Assembly of Mitochondrial Cardiolipins. Cell Rep. 30: 4281–4291.e4.

16. Pennington, E. R., K. Funai, D. A. Brown, and S. R. Shaikh. 2019. The role of cardiolipin concentration and acyl chain composition on mitochondrial inner membrane molecular organization and function. Biochim. Biophys. Acta Mol. Cell Biol. Lipids. 1864: 1039–1052.

17. Schlame, M.. 2013. Cardiolipin remodeling and the function of tafazzin. Biochim. Biophys. Acta. 1831: 582–588.

18. Ye, C., Z. Shen, and M. L. Greenberg. 2016. Cardiolipin remodeling: a regulatory hub for modulating cardiolipin metabolism and function. J. Bioenerg. Biomembr. 48: 113–123.

19. Xu, Y., M. Anjaneyulu, A. Donelian, W. Yu, M. L. Greenberg, M. Ren, E. Owusu-Ansah, and M. Schlame. 2019. Assembly of the complexes of oxidative phosphorylation triggers the remodeling of cardiolipin. Proc. Natl. Acad. Sci. U. S. A. 116: 11235–11240.

20. Xu, Y., H. Erdjument-Bromage, C. K. L. Phoon, T. A. Neubert, M. Ren, and M. Schlame. 2021. Cardiolipin remodeling enables protein crowding in the inner mitochondrial membrane. EMBO J. 40: e108428.

21. Malhotra, A., I. Edelman-Novemsky, Y. Xu, H. Plesken, J. Ma, M. Schlame, and M. Ren. 2009. Role of calcium-independent phospholipase A2 in the pathogenesis of Barth syndrome. Proc. Natl. Acad. Sci. U. S. A. 106: 2337–2341.

22. Malhotra, A., Y. Xu, M. Ren, and M. Schlame. 2009. Formation of molecular species of mitochondrial cardiolipin. 1. A novel transacylation mechanism to shuttle fatty acids between sn-1 and sn-2 positions of multiple phospholipid species. Biochim. Biophys. Acta. 1791: 314–320.

23. Schlame, M., D. Acehan, B. Berno, Y. Xu, S. Valvo, M. Ren, D. L. Stokes, and R. M. Epand. 2012. The physical state of lipid substrates provides transacylation specificity for tafazzin. Nat. Chem. Biol. 8: 862–869.

24. Claypool, S. M., and C. M. Koehler. 2012. The complexity of cardiolipin in health and disease. Trends in Biochemical Sciences. 37: 32–41. [online] 10.1016/j.tibs.2011.09.003.

25. Falabella, M., H. J. Vernon, M. G. Hanna, S. M. Claypool, and R. D. S. Pitceathly. 2021. Cardiolipin, Mitochondria, and Neurological Disease. Trends Endocrinol. Metab. 32: 224–237.

26. Adès, L. C., A. K. Gedeon, M. J. Wilson, M. Latham, M. W. Partington, J. C. Mulley, J. Nelson, K. Lui, and D. O. Sillence. 1993. Barth syndrome: Clinical features and confirmation of gene localisation to distal Xq28. American Journal of Medical Genetics. 45: 327–334. [online] 10.1002/ajmg.1320450309.

27. Bione, S., P. D’Adamo, E. Maestrini, A. K. Gedeon, P. A. Bolhuis, and D. Toniolo. 1996. A novel X-linked gene, G4.5. is responsible for Barth syndrome. Nat. Genet. 12: 385–389.

28. Vreken, P., F. Valianpour, L. G. Nijtmans, L. A. Grivell, B. Plecko, R. J. Wanders, and P. G. Barth. 2000. Defective remodeling of cardiolipin and phosphatidylglycerol in Barth syndrome. Biochem. Biophys. Res. Commun. 279: 378–382.

29. Schlame, M., J. A. Towbin, P. M. Heerdt, R. Jehle, S. DiMauro, and T. J. J. Blanck. 2002. Deficiency of tetralinoleoyl-cardiolipin in Barth syndrome. Ann. Neurol. 51: 634–637.

30. van Werkhoven, M. A., D. R. Thorburn, A. K. Gedeon, and J. J. Pitt. 2006. Monolysocardiolipin in cultured fibroblasts is a sensitive and specific marker for Barth Syndrome. J. Lipid Res. 47: 2346–2351.

31. Byeon, S. K., M. G. Ramarajan, A. K. Madugundu, D. Oglesbee, H. J. Vernon, and A. Pandey. 2021. High-resolution mass spectrometric analysis of cardiolipin profiles in Barth syndrome. Mitochondrion. 60: 27–32.

32. Barth, P. G., H. R. Scholte, J. A. Berden, J. M. Van der Klei-Van Moorsel, I. E. Luyt-Houwen, E. T. Van’t Veer-Korthof, J. J. Van der Harten, and M. A. Sobotka-Plojhar. 1983. An X-linked mitochondrial disease affecting cardiac muscle, skeletal muscle and neutrophil leucocytes. J. Neurol. Sci. 62: 327–355.

33. Raja, V., C. A. Reynolds, and M. L. Greenberg. 2017. Barth syndrome: A life-threatening disorder caused by abnormal cardiolipin remodeling. J Rare Dis Res Treat. 2: 58–62.

34. Claypool, S. M., J. M. McCaffery, and C. M. Koehler. 2006. Mitochondrial mislocalization and altered assembly of a cluster of Barth syndrome mutant tafazzins. J. Cell Biol. 174: 379–390.

35. Claypool, S. M., K. Whited, S. Srijumnong, X. Han, and C. M. Koehler. 2011. Barth syndrome mutations that cause tafazzin complex lability. J. Cell Biol. 192: 447–462.

36. Xu, Y., C. K. L. Phoon, B. Berno, K. D’Souza, E. Hoedt, G. Zhang, T. A. Neubert, R. M. Epand, M. Ren, and M. Schlame. 2016. Loss of protein association causes cardiolipin degradation in Barth syndrome. Nat. Chem. Biol. 12: 641–647.

37. Houtkooper, R. H., R. J. Rodenburg, C. Thiels, H. van Lenthe, F. Stet, B. T. Poll-The, J. E. Stone, C. G. Steward, R. J. Wanders, J. Smeitink, W. Kulik, and F. M. Vaz. 2009. Cardiolipin and monolysocardiolipin analysis in fibroblasts, lymphocytes, and tissues using high-performance liquid chromatography-mass spectrometry as a diagnostic test for Barth syndrome. Anal. Biochem. 387: 230–237.

38. Ackermann, E. J., K. Conde-Frieboes, and E. A. Dennis. 1995. Inhibition of macrophage Ca(2+)-independent phospholipase A2 by bromoenol lactone and trifluoromethyl ketones. J. Biol. Chem. 270: 445–450.

39. Anzmann, A. F., O. L. Sniezek, A. Pado, V. Busa, F. M. Vaz, S. D. Kreimer, L. R. DeVine, R. N. Cole, A. Le, B. J. Kirsch, S. M. Claypool, and H. J. Vernon. 2021. Diverse mitochondrial abnormalities in a new cellular model of TAFFAZZIN deficiency are remediated by cardiolipin-interacting small molecules. J. Biol. Chem. 297. [online] https://www.jbc.org/article/S0021-9258(21)00807-3/fulltext (Accessed April 7, 2024).

40. Wang, G., M. L. McCain, L. Yang, A. He, F. S. Pasqualini, A. Agarwal, H. Yuan, D. Jiang, D. Zhang, L. Zangi, J. Geva, A. E. Roberts, Q. Ma, J. Ding, J. Chen, D.-Z. Wang, K. Li, J. Wang, R. J. A. Wanders, W. Kulik, F. M. Vaz, M. A. Laflamme, C. E. Murry, K. R. Chien, R. I. Kelley, G. M. Church, K. K. Parker, and W. T. Pu. 2014. Modeling the mitochondrial cardiomyopathy of Barth syndrome with induced pluripotent stem cell and heart-on-chip technologies. Nat. Med. 20: 616–623.

41. Ji, J., and M. L. Greenberg. 2022. Cardiolipin function in the yeast S. cerevisiae and the lessons learned for Barth syndrome. J. Inherit. Metab. Dis. 45: 60–71.

42. Pu, W. T.. 2022. Experimental models of Barth syndrome. J. Inherit. Metab. Dis. 45: 72–81.

43. Beranek, A., G. Rechberger, H. Knauer, H. Wolinski, S. D. Kohlwein, and R. Leber. 2009. Identification of a cardiolipin-specific phospholipase encoded by the gene CLD1 (YGR110W) in yeast. J. Biol. Chem. 284: 11572–11578.

44. Abe, M., Y. Hasegawa, M. Oku, Y. Sawada, E. Tanaka, Y. Sakai, and H. Miyoshi. 2016. Mechanism for Remodeling of the Acyl Chain Composition of Cardiolipin Catalyzed by Saccharomyces cerevisiae Tafazzin. J. Biol. Chem. 291: 15491–15502.

45. Schlame, M., Y. Xu, and M. Ren. 2017. The Basis for Acyl Specificity in the Tafazzin Reaction. J. Biol. Chem. 292: 5499–5506.

46. Gu, Z., F. Valianpour, S. Chen, F. M. Vaz, G. A. Hakkaart, R. J. A. Wanders, and M. L. Greenberg. 2004. Aberrant cardiolipin metabolism in the yeast taz1 mutant: a model for Barth syndrome. Mol. Microbiol. 51: 149–158.

47. Baile, M. G., M. Sathappa, Y.-W. Lu, E. Pryce, K. Whited, J. Michael McCaffery, X. Han, N. N. Alder, and S. M. Claypool. 2014. Unremodeled and Remodeled Cardiolipin Are Functionally Indistinguishable in Yeast. Journal of Biological Chemistry. 289: 1768–1778. [online] 10.1074/jbc.m113.525733.

48. Brandner, K., D. U. Mick, A. E. Frazier, R. D. Taylor, C. Meisinger, and P. Rehling. 2005. Taz1, an outer mitochondrial membrane protein, affects stability and assembly of inner membrane protein complexes: implications for Barth Syndrome. Mol. Biol. Cell. 16: 5202–5214.

49. Ye, C., W. Lou, Y. Li, I. A. Chatzispyrou, M. Hüttemann, I. Lee, R. H. Houtkooper, F. M. Vaz, S. Chen, and M. L. Greenberg. 2014. Deletion of the cardiolipin-specific phospholipase Cld1 rescues growth and life span defects in the tafazzin mutant: implications for Barth syndrome. J. Biol. Chem. 289: 3114–3125.

50. Chen, S., Q. He, and M. L. Greenberg. 2008. Loss of tafazzin in yeast leads to increased oxidative stress during respiratory growth. Mol. Microbiol. 68: 1061–1072.

51. Ma, L., F. M. Vaz, Z. Gu, R. J. A. Wanders, and M. L. Greenberg. 2004. The human TAZ gene complements mitochondrial dysfunction in the yeast taz1Delta mutant. Implications for Barth syndrome. J. Biol. Chem. 279: 44394–44399.

52. Jiang, F., M. T. Ryan, M. Schlame, M. Zhao, Z. Gu, M. Klingenberg, N. Pfanner, and M. L. Greenberg. 2000. Absence of cardiolipin in the crd1 null mutant results in decreased mitochondrial membrane potential and reduced mitochondrial function. J. Biol. Chem. 275: 22387–22394.

53. Baile, M. G., K. Whited, and S. M. Claypool. 2013. Deacylation on the matrix side of the mitochondrial inner membrane regulates cardiolipin remodeling. Mol. Biol. Cell. 24: 2008–2020.

54. Venkatraman, K., C. T. Lee, G. C. Garcia, A. Mahapatra, D. Milshteyn, G. Perkins, K.-Y. Kim, H. A. Pasolli, S. Phan, J. Lippincott-Schwartz, M. H. Ellisman, P. Rangamani, and I. Budin. 2023. Cristae formation is a mechanical buckling event controlled by the inner mitochondrial membrane lipidome. EMBO J. e114054.

55. Kwast, K. E., P. V. Burke, B. T. Staahl, and R. O. Poyton. 1999. Oxygen sensing in yeast: Evidence for the involvement of the respiratory chain in regulating the transcription of a subset of hypoxic genes. Proceedings of the National Academy of Sciences. 96: 5446–5451. [online] 10.1073/pnas.96.10.5446.

56. Vasconcelles, M. J., Y. Jiang, K. McDaid, L. Gilooly, S. Wretzel, D. L. Porter, C. E. Martin, and M. A. Goldberg. 2001. Identification and Characterization of a Low Oxygen Response Element Involved in the Hypoxic Induction of a Family ofSaccharomyces cerevisiae Genes. Journal of Biological Chemistry. 276: 14374–14384. [online] 10.1074/jbc.m009546200.

57. Wong, A. M., and I. Budin. 2024. Organelle-targeted Laurdans measure heterogeneity in subcellular membranes and their responses to saturated lipid stress. bioRxiv. [online] 10.1101/2024.04.16.589828.

58. Ståhlman, M., C. S. Ejsing, K. Tarasov, J. Perman, J. Borén, and K. Ekroos. 2009. High-throughput shotgun lipidomics by quadrupole time-of-flight mass spectrometry. Journal of Chromatography B. 877: 2664–2672. [online] 10.1016/j.jchromb.2009.02.037.

59. Klose, C., M. A. Surma, M. J. Gerl, F. Meyenhofer, A. Shevchenko, and K. Simons. 2012. Flexibility of a Eukaryotic Lipidome – Insights from Yeast Lipidomics. PLoS ONE. 7: e35063. [online] 10.1371/journal.pone.0035063.

60. Ejsing, C. S., J. L. Sampaio, V. Surendranath, E. Duchoslav, K. Ekroos, R. W. Klemm, K. Simons, and A. Shevchenko. 2009. Global analysis of the yeast lipidome by quantitative shotgun mass spectrometry. Proceedings of the National Academy of Sciences. 106: 2136–2141. [online] 10.1073/pnas.0811700106.

61. Surma, M. A., R. Herzog, A. Vasilj, C. Klose, N. Christinat, D. Morin-Rivron, K. Simons, M. Masoodi, and J. L. Sampaio. 2015. An automated shotgun lipidomics platform for high throughput, comprehensive, and quantitative analysis of blood plasma intact lipids. European Journal of Lipid Science and Technology. 117: 1540–1549. [online] 10.1002/ejlt.201500145.

62. Herzog, R., D. Schwudke, K. Schuhmann, J. L. Sampaio, S. R. Bornstein, M. Schroeder, and A. Shevchenko. 2011. A novel informatics concept for high-throughput shotgun lipidomics based on the molecular fragmentation query language. Genome Biol. 12: R8.

63. Herzog, R., K. Schuhmann, D. Schwudke, J. L. Sampaio, S. R. Bornstein, M. Schroeder, and A. Shevchenko. 2012. LipidXplorer: a software for consensual cross-platform lipidomics. PLoS One. 7: e29851.

64. Fuhrmann, D. C., and B. Brüne. 2017. Mitochondrial composition and function under the control of hypoxia. Redox Biol. 12: 208–215.

65. Klecker, T., and B. Westermann. 2021. Pathways shaping the mitochondrial inner membrane. Open Biol. 11: 210238.

66. Degreif, D., T. de Rond, A. Bertl, J. D. Keasling, and I. Budin. 2017. Lipid engineering reveals regulatory roles for membrane fluidity in yeast flocculation and oxygen-limited growth. Metab. Eng. 41: 46–56.

67. Kamphorst, J. J., J. R. Cross, J. Fan, E. de Stanchina, R. Mathew, E. P. White, C. B. Thompson, and J. D. Rabinowitz. 2013. Hypoxic and Ras-transformed cells support growth by scavenging unsaturated fatty acids from lysophospholipids. Proc. Natl. Acad. Sci. U. S. A. 110: 8882–8887.

68. Ackerman, D., S. Tumanov, B. Qiu, E. Michalopoulou, M. Spata, A. Azzam, H. Xie, M. C. Simon, and J. J. Kamphorst. 2018. Triglycerides Promote Lipid Homeostasis during Hypoxic Stress by Balancing Fatty Acid Saturation. Cell Rep. 24: 2596–2605.e5.

69. Mthembu, S. X. H., S. E. Mazibuko-Mbeje, S. Silvestri, P. Orlando, F. Marcheggiani, I. Cirilli, B. B. Nkambule, C. J. F. Muller, L. Tiano, and P. V. Dludla. 2024. Low levels and partial exposure to palmitic acid improves mitochondrial function and the oxidative status of cultured cardiomyoblasts. Toxicol Rep. 12: 234–243.

70. Hsu, P., X. Liu, J. Zhang, H.-G. Wang, J.-M. Ye, and Y. Shi. 2015. Cardiolipin remodeling by TAZ/tafazzin is selectively required for the initiation of mitophagy. Autophagy. 11: 643–652.

71. Shen, Z., Y. Li, A. N. Gasparski, H. Abeliovich, and M. L. Greenberg. 2017. Cardiolipin Regulates Mitophagy through the Protein Kinase C Pathway *. J. Biol. Chem. 292: 2916–2923.

72. Sakakibara, K., A. Eiyama, S. W. Suzuki, M. Sakoh-Nakatogawa, N. Okumura, M. Tani, A. Hashimoto, S. Nagumo, N. Kondo-Okamoto, C. Kondo-Kakuta, E. Asai, H. Kirisako, H. Nakatogawa, O. Kuge, T. Takao, Y. Ohsumi, and K. Okamoto. 2015. Phospholipid methylation controls Atg32-mediated mitophagy and Atg8 recycling. EMBO J. 34: 2703–2719–2719.

73. Kanki, T., K. Wang, Y. Cao, M. Baba, and D. J. Klionsky. 2009. Atg32 is a mitochondrial protein that confers selectivity during mitophagy. Dev. Cell. 17: 98–109.

74. Urafuji, K., and M. Arioka. 2016. Yor022c protein is a phospholipase A1 that localizes to the mitochondrial matrix. Biochem. Biophys. Res. Commun. 480: 302–308.

75. Yadav, P. K., and R. Rajasekharan. 2016. Misregulation of a DDHD Domain-containing Lipase Causes Mitochondrial Dysfunction in Yeast. J. Biol. Chem. 291: 18562–18581.

76. Rand, R. P., and S. Sengupta. 1972. Cardiolipin forms hexagonal structures with divalent cations. Biochim. Biophys. Acta. 255: 484–492.

77. Cullis, P. R., A. J. Verkleij, and P. H. Ververgaert. 1978. Polymorphic phase behaviour of cardiolipin as detected by 31P NMR and freeze-fracture techniques. Effects of calcium, dibucaine and chlorpromazine. Biochim. Biophys. Acta. 513: 11–20.

78. Lewis, R. N. A. H., and R. N. McElhaney. 2009. The physicochemical properties of cardiolipin bilayers and cardiolipin-containing lipid membranes. Biochimica et Biophysica Acta (BBA) – Biomembranes. 1788: 2069–2079.

79. Lewis, R. N. A. H., D. Zweytick, G. Pabst, K. Lohner, and R. N. McElhaney. 2007. Calorimetric, x-ray diffraction, and spectroscopic studies of the thermotropic phase behavior and organization of tetramyristoyl cardiolipin membranes. Biophys. J. 92: 3166–3177.

80. Barbot, M., D. C. Jans, C. Schulz, N. Denkert, B. Kroppen, M. Hoppert, S. Jakobs, and M. Meinecke. 2015. Mic10 oligomerizes to bend mitochondrial inner membranes at cristae junctions. Cell Metab. 21: 756–763.

81. Tarasenko, D., M. Barbot, D. C. Jans, B. Kroppen, B. Sadowski, G. Heim, W. Möbius, S. Jakobs, and M. Meinecke. 2017. The MICOS component Mic60 displays a conserved membrane-bending activity that is necessary for normal cristae morphology. J. Cell Biol. 216: 889–899.

82. Blum, T. B., A. Hahn, T. Meier, K. M. Davies, and W. Kühlbrandt. 2019. Dimers of mitochondrial ATP synthase induce membrane curvature and self-assemble into rows. Proc. Natl. Acad. Sci. U. S. A. 116: 4250–4255.

83. Powell, G. L., and D. Marsh. 1985. Polymorphic phase behavior of cardiolipin derivatives studied by phosphorus-31 NMR and x-ray diffraction. Biochemistry. 24: 2902–2908. [online] 10.1021/bi00333a013.

84. Boyd, K. J., N. N. Alder, and E. R. May. 2018. Molecular Dynamics Analysis of Cardiolipin and Monolysocardiolipin on Bilayer Properties. Biophys. J. 114: 2116–2127.

85. Duncan, A. L.. 2020. Monolysocardiolipin (MLCL) interactions with mitochondrial membrane proteins. Biochem. Soc. Trans. 48: 993–1004.

86. Valianpour, F., V. Mitsakos, D. Schlemmer, J. A. Towbin, J. M. Taylor, P. G. Ekert, D. R. Thorburn, A. Munnich, R. J. A. Wanders, P. G. Barth, and F. M. Vaz. 2005. Monolysocardiolipins accumulate in Barth syndrome but do not lead to enhanced apoptosis. J. Lipid Res. 46: 1182–1195.

87. Saric, A., K. Andreau, A.-S. Armand, I. M. Møller, and P. X. Petit. 2015. Barth Syndrome: From Mitochondrial Dysfunctions Associated with Aberrant Production of Reactive Oxygen Species to Pluripotent Stem Cell Studies. Front. Genet. 6: 359.

88. Oemer, G., M.-L. Edenhofer, Y. Wohlfarter, K. Lackner, G. Leman, J. Koch, L. H. D. Cardoso, H. H. Lindner, E. Gnaiger, S. Dubrac, J. Zschocke, and M. A. Keller. 2021. Fatty acyl availability modulates cardiolipin composition and alters mitochondrial function in HeLa cells. J. Lipid Res. 62: 100111.

89. Oemer, G., J. Koch, Y. Wohlfarter, K. Lackner, R. E. M. Gebert, S. Geley, J. Zschocke, and M. A. Keller. 2022. The lipid environment modulates cardiolipin and phospholipid constitution in wild type and tafazzin-deficient cells. J. Inherit. Metab. Dis. 45: 38–50.

90. Zhu, S., J. Pang, A. Nguyen, H. Huynh, S. Lee, Y. Gu, F. M. Vaz, and X. Fang. 2024. Dietary linoleic acid supplementation fails to rescue established cardiomyopathy in Barth syndrome. Journal of Molecular and Cellular Cardiology Plus. 8: 100076.

91. Ostrander, D. B., G. C. Sparagna, A. A. Amoscato, J. B. McMillin, and W. Dowhan. 2001. Decreased cardiolipin synthesis corresponds with cytochrome c release in palmitate-induced cardiomyocyte apoptosis. J. Biol. Chem. 276: 38061–38067.

92. Higgs, H. N., and J. A. Glomset. 1996. Purification and properties of a phosphatidic acid-preferring phospholipase A1 from bovine testis. Examination of the molecular basis of its activation. J. Biol. Chem. 271: 10874–10883.

93. Tesson, C., M. Nawara, M. A. M. Salih, R. Rossignol, M. S. Zaki, M. Al Balwi, R. Schule, C. Mignot, E. Obre, A. Bouhouche, F. M. Santorelli, C. M. Durand, A. C. Oteyza, K. H. El-Hachimi, A. Al Drees, N. Bouslam, F. Lamari, S. A. Elmalik, M. M. Kabiraj, M. Z. Seidahmed, T. Esteves, M. Gaussen, M.-L. Monin, G. Gyapay, D. Lechner, M. Gonzalez, C. Depienne, F. Mochel, J. Lavie, L. Schols, D. Lacombe, M. Yahyaoui, I. Al Abdulkareem, S. Zuchner, A. Yamashita, A. Benomar, C. Goizet, A. Durr, J. G. Gleeson, F. Darios, A. Brice, and G. Stevanin. 2012. Alteration of fatty-acid-metabolizing enzymes affects mitochondrial form and function in hereditary spastic paraplegia. Am. J. Hum. Genet. 91: 1051–1064.

94. Yamaguchi-Iwai, Y., A. Dancis, and R. D. Klausner. 1995. AFT1: a mediator of iron regulated transcriptional control in Saccharomyces cerevisiae. EMBO J. 14: 1231–1239–1239.

95. Clarke, S. L. N., A. Bowron, I. L. Gonzalez, S. J. Groves, R. Newbury-Ecob, N. Clayton, R. P. Martin, B. Tsai-Goodman, V. Garratt, M. Ashworth, V. M. Bowen, K. R. McCurdy, M. K. Damin, C. T. Spencer, M. J. Toth, R. I. Kelley, and C. G. Steward. 2013. Barth syndrome. Orphanet J. Rare Dis. 8: 23.

